# Thalamic feedback shapes brain responses evoked by cortical stimulation in mice and humans

**DOI:** 10.1101/2024.01.31.578243

**Authors:** Simone Russo, Leslie Claar, Lydia Marks, Giri Krishnan, Giulia Furregoni, Flavia Maria Zauli, Gabriel Hassan, Michela Solbiati, Piergiorgio d’Orio, Ezequiel Mikulan, Simone Sarasso, Mario Rosanova, Ivana Sartori, Maxim Bazhenov, Andrea Pigorini, Marcello Massimini, Christof Koch, Irene Rembado

## Abstract

Cortical stimulation with single pulses is a common technique in clinical practice and research. However, we still do not understand the extent to which it engages subcortical circuits which contribute to the associated evoked potentials (EPs). Here we find that cortical stimulation generates remarkably similar EPs in humans and mice, with a late component similarly modulated by the subject’s behavioral state. We optogenetically dissect the underlying circuit in mice, demonstrating that the late component of these EPs is caused by a thalamic hyperpolarization and rebound. The magnitude of this late component correlates with the bursting frequency and synchronicity of thalamic neurons, modulated by the subject’s behavioral state. A simulation of the thalamo-cortical circuit highlights that both intrinsic thalamic currents as well as cortical and thalamic GABAergic neurons contribute to this response profile. We conclude that the cortical stimulation engages cortico-thalamo-cortical circuits highly preserved across different species and stimulation modalities.

**Graphical abstract:** 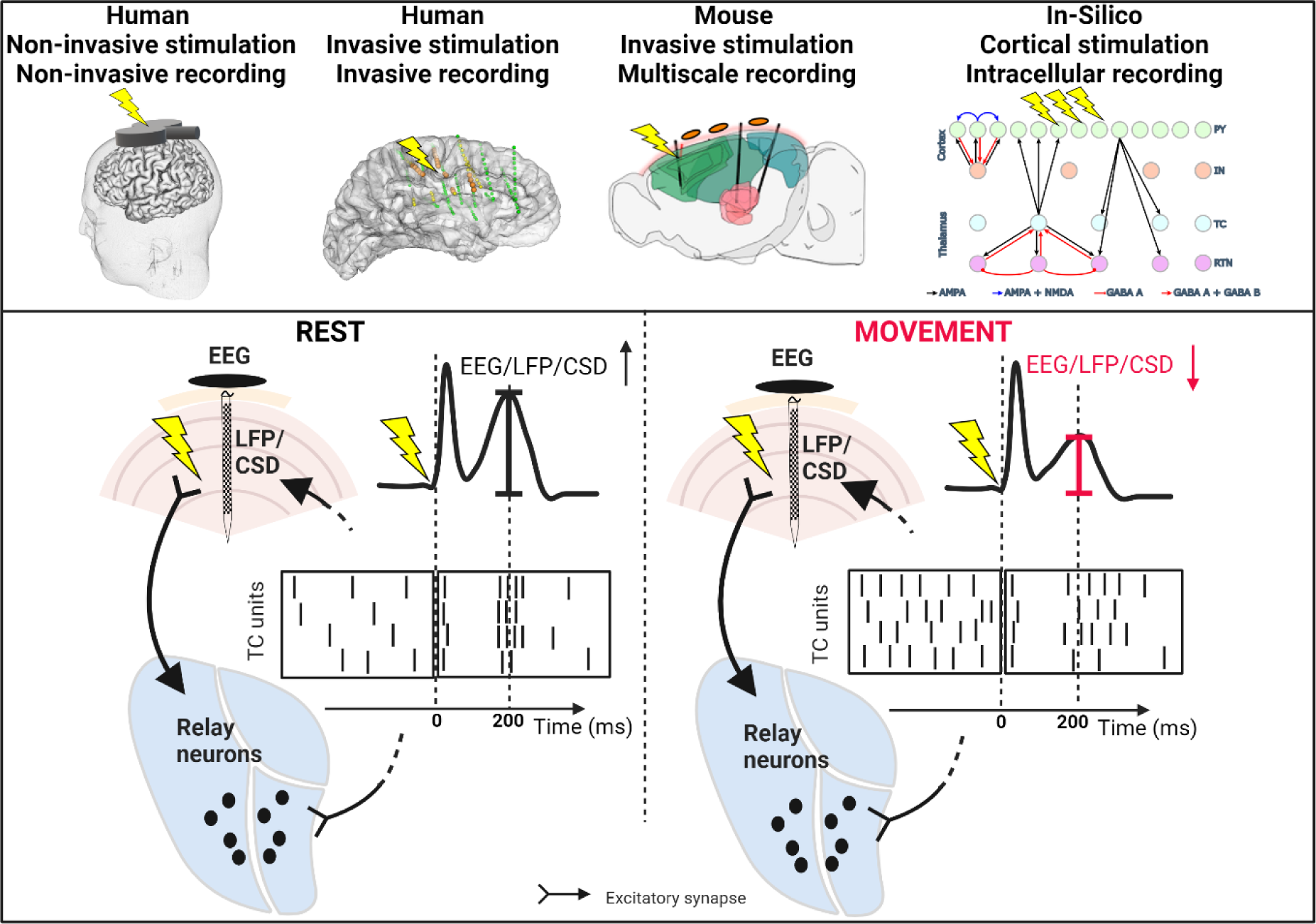

## Introduction

Evoked potentials (**EPs**), inferred from electroencephalography (**EEG**) by means of trial averaging, are a common tool to investigate the relationship between neural activity, human cognition, and clinical disorders^1–3^. EPs provide a non-invasive and direct measurement of neural activity with high temporal resolution. The EPs elicited by cortical stimulation – either delivered invasively through electrical stimulation (**ES**), or non-invasively via transcranial magnetic stimulation (**TMS**) – are commonly used for functional mapping of brain networks^4–7^, the diagnosis of brain dysfunctions^8–10^, and the detection of consciousness in behaviorally unresponsive patients^11–16^. Moreover, EPs elicited by cortical stimulation speak directly to a central conundrum of systems neuroscience: to what extent does neural activity propagate across areas via cortico-cortical and/or cortico-subcortico-cortical projections? Understanding the contribution of feedforward and of feedback projections to EPs is important to properly interpret experimental and clinical results.

TMS non-invasively stimulates the brain by electromagnetically inducing current flow in the cortical tissue underneath the TMS coil resting against the scalp. Despite its widespread use over 30 years of research and clinical practice^17–19^, the responses evoked by TMS have been a source of debate due to the potential contribution of peripherally induced somatosensory and auditory components of the TMS stimulation^20^. Against this hypothesis, single neurons recordings in awake non-human primates demonstrated that TMS can induce the same stereotyped triphasic spiking pattern evoked by ES and previously reported in different species^21–26^. These responses to ES are intrinsically free of peripheral confounds and can reverberate through the cortical network for several hundreds of milliseconds after stimulus delivery^27^. Along the same lines, a more recent study in humans showed that TMS-EPs can propagate in the network for hundreds of milliseconds, and that such responses highly resemble the EPs generated by delivering ES to the same area^28^. These findings support the notion that TMS responses are not due to peripheral activations, but rather due to recurrent interactions across different regions^29^. Indeed, ES directly excites nearby neurons that cause secondary responses in connected neuronal populations^7,30^. Responses to cortical ES are commonly used to define the connectivity across cortical areas, to the point that they are also known as cortico-cortical evoked potentials^4–7^, implicitly ruling out contributions from subcortical structures. In contrast to this perspective, in our previous study, we proposed that cortico-thalamo-cortical interactions drive long-lasting EEG-EPs^21^ in mice. Indeed, thalamocortical neurons receive major inputs from the neocortex and in turn their outputs project back to it^31–33^. As such, thalamocortical neurons are strategically positioned to relay information between cortical areas and they could be considered a major generator of the cortical postsynaptic potentials captured by the EEG^34,35^ which contribute to the EP observed after cortical stimulation. Given the challenge of recording thalamic single neurons in humans, any thalamic contributions to EPs can be only indirectly inferred^36^ and remain largely unexplored.

Here we show that, despite the differences between species (humans versus mice) and between magnetic and electrical pulses, the late responses evoked by cortical stimulation are remarkably similar and likewise modulated by the behavioral state of the subject, defined as either movement or resting state. The fact that responses to cortical stimulation are invariably modulated by movement suggests that a common neuronal mechanism underlies these responses. By combining causal manipulation via optogenetic intervention with multi-scale recordings (single units, LFP and EEG) in mice, and modeling of the cortico-thalamic circuits, we demonstrate that the late component in the EPs following direct cortical stimulation depends on the thalamus. It is therefore likely that a common mechanism, involving thalamic intrinsic currents and cortical and thalamic GABAergic neurons, shapes the late responses to cortical stimulation in both species. In this perspective, the present study addresses fundamental questions around cortical stimulation by providing compelling evidence that the long-lasting EEG responses to both TMS and ES in humans reflect cortico-thalamo-cortical interactions, are modulated by the behavioral state of the subject, and reveal key information about the state of the thalamocortical network.

## Results

### Multiscale late EP responses elicited by cortical stimulation are modulated by behavioral state in humans and mice

We start by describing the basic cortical EP in humans and its modulation by behavioral state. EEG responses evoked by single pulse TMS delivered over the left premotor cortex were recorded in a group of 12 healthy volunteers (for demographics see Table S1) during rest and movement. For the baseline rest condition, subjects were asked to stay awake while relaxing their hands; for the movement condition, subjects were asked to intermittently squeeze a rubber ball with the right hand. For these two conditions, we evaluated the early EP component evoked by TMS in the time window 3-50 ms from stimulation onset (usually ascribed to monosynaptic pathways) and the late response in the 150-250 ms window (usually ascribed to polysynaptic pathways; Figure 1A).

**Figure 1.**
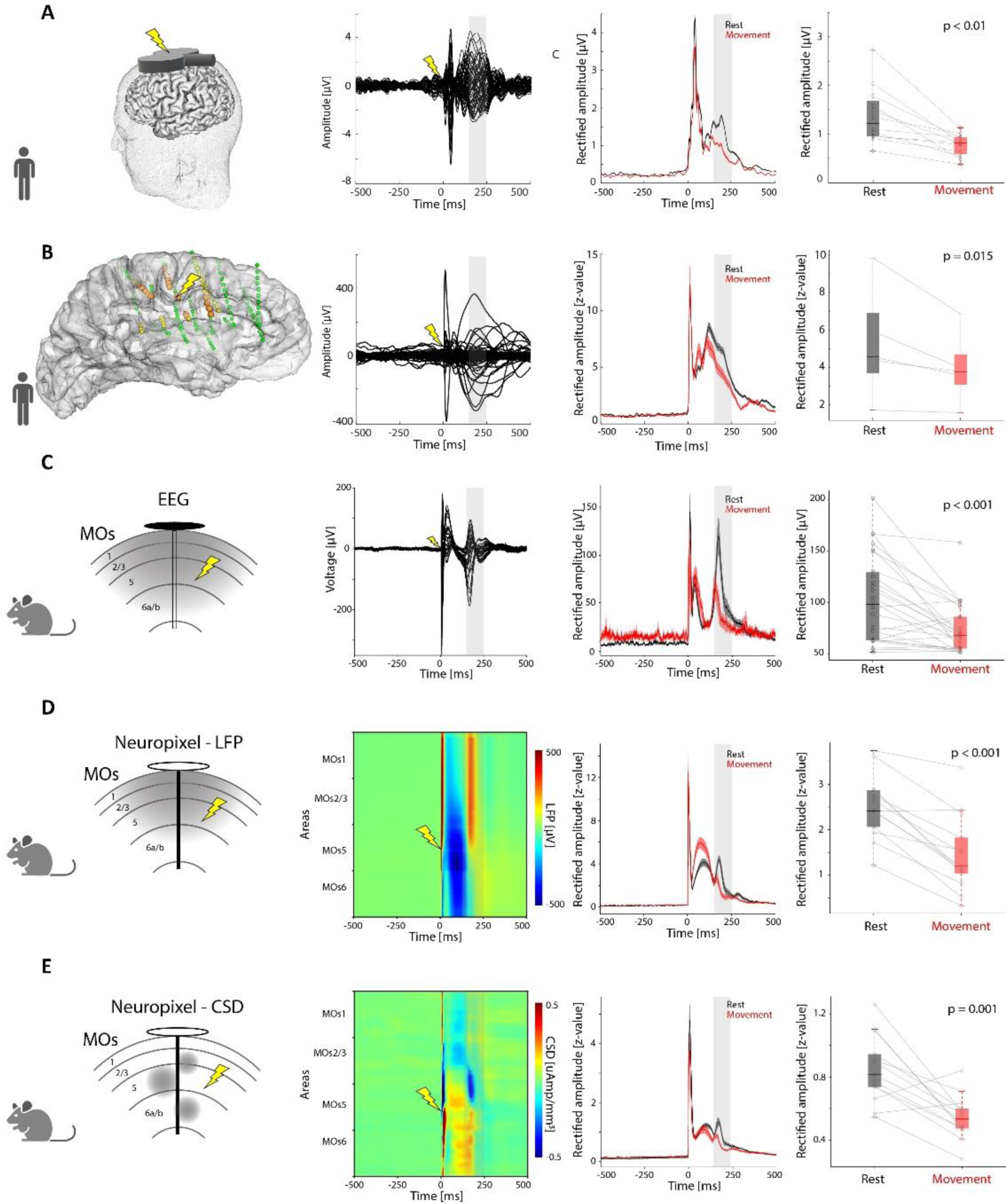
Behavioral-state-dependent modulation of cortico-thalamo-cortical responses evoked by the electrical and magnetic stimulation of MOs in mice and premotor cortex in humans. A) Modulation of the EEG responses to TMS delivered to the premotor cortex in humans at rest and during movement. Left to right: Schematic of the setup with a TMS coil stimulating the premotor area. Butterfly plot of TMS-evoked responses (-500 to 500 ms around stimulus onset) for all contact traces of one representative subject. Grand-average (12 subjects) during rest (black) and movement (red). The rectified amplitude average of the late component (150-250 ms; shaded gray) during rest and movement for each subject (Wilcoxon signed rank test: p<0.01; same subjects are connected by lines). B) Modulation of the iEEG responses to ES in humans at rest and during movement. Left to right: Schematic of the setup with 3D reconstruction from one representative subject of the stimulated cortical area (lightning symbol), the recording contacts within the region of interest (ROI) responding (orange dots) and non-responding (yellow dots) to the stimulation, and the contacts outside the ROI (green dots). Butterfly plot of single pulse ES-evoked responses (-500 to 500 ms around stimulus onset) for all contacts from one representative subject. Evoked iEEG rectified amplitude during rest (black) and movement (red) from 105 contacts and 5 subjects. Intracerebral EEG rectified amplitude average over the late response window (150-250 ms, shaded gray) at rest and movement for all responding ROI contacts across subjects (Linear Mixed Effect Model: p=0.015; same subjects are connected by lines). C) Modulation of the EEG responses evoked by the strong (66.15±12.11 uA) electrical stimulation of the deep layers in MOs at rest and during movement. Left to right: representation of the area recorded by an EEG electrode (grey) and the stimulated site (lightning symbol). Butterfly plot of the EPs (-500 to 500 ms around stimulus onset) for all 30 EEG electrode traces averaged across subjects. EEG rectified amplitude evoked at rest (black) and movement (red) calculated over all EEG contacts for all subjects (26 sessions, 15 subjects). Averaged EEG rectified amplitude over the late response window (150-250 ms, shaded gray) at rest and movement for all EEG electrodes across subjects (Wilcoxon signed rank test p<0.001; same subjects are connected by lines). D) Modulation of the LFP responses from the Neuropixels probe in MOs, averaged over n=13 subjects, evoked by the electrical stimulation of the deep layers of the same area (lightning symbol) at rest and during movement. Plots are structured in the same fashion as in C, with the voltages displayed in a color map as function of depth from superficial to deep layers (represented in y-axis from top to bottom). The amplitude over the late response window (150-250 ms, shaded gray) is larger at rest compared to movement (Wilcoxon signed rank test; Rectified amplitude: p<0. 001). E) Modulation of the CSD, averaged over n=13 subjects, estimated from the Neuropixels probe in MOs where the electrical stimulation was delivered, at rest and during movement. Same structure as D. The amplitude over the late response window (150-250 ms, shaded gray) is larger at rest compared to movement (Wilcoxon signed rank test; p=0.001).

Compared to rest, both the rectified amplitude of the late response over premotor areas (Fc1, Fcz, Fc2, C1, Cz, C2), and its associated phase locking factor (PLF, a measure of the determinism of the response across trials^37^) were significantly reduced during movement (amplitude at rest: 1.37±0.59 μV versus during movement: 0.77±0.22 μV p<0.01 Figure 1A; PLF at rest 0.21±0.07 versus during movement 0.15±0.05; p=0.002, Figure S1A). Similarly, a passive movement of the same hand induced a significant modulation of the late component (Figure S2A). Importantly, the response modulation was not caused by potential somatosensory confounds related to TMS, as shown by a lack of a similar modulation in the somatosensory potentials (SSEPs) evoked by electrically stimulating the left median nerve (Figure S2D). We also observed adaptation mechanisms specific for SSEP, but not for the TMS-EEG responses (Figure S2E), corroborating the hypothesis that the late components evoked by TMS and sensory stimulation are most likely generated by different neuronal circuits. Moreover, the observed movement-related modulation was network specific, as shown in a subset of 4 subjects (5 sessions) whose evoked responses from the same contacts above the premotor area elicited by TMS delivered to posterior areas (parietal and occipital lobes) showed no significant modulation in either active or passive movement compared to rest (Figure S2C). As opposed to the late component, the early component of the EP (3-50 ms from stimulus onset) showed a reduction in amplitude, but not in phase locking (amplitude at rest 2.24±0.45 μV versus 1.88±0.32 μV during movement; PLF at rest: 0.43±0.8 versus movement: 0.43±0.9; amplitude: p<0.01; PLF: p=0.97; Figure S2A), although the decreases in rectified amplitude of the early and late components were not significantly correlated (Figure S2B; Spearman’s Correlation; p=0.13; r=0.46). These results show that the amplitude of the TMS-evoked EP in the 150-250 ms window is reduced during active and passive movement of the hand, an effect that is network specific and not related to somatosensory stimulation.

A similar and significant modulation of the late responses was observed in patients with epilepsy implanted with intracerebral EEG (iEEG) electrodes for presurgical screening (5 subjects; for demographics see Table S2; Figure 1B). In this case, the single pulse ES were delivered invasively between two adjacent contacts within one iEEG depth shaft in the contralateral premotor area of the hand performing the task. The late responses evoked in nearby iEEG contacts (within 3 cm from the stimulated site) showed a significant reduction of the evoked amplitude (Figure 1B, z-scored amplitude 5.95±5.79 at rest versus 4.30±4.18 z-value during movement; p<0.05) and PLF during movement (Figure S1A, PLF during rest 0.55±0.27 versus 0.49±0.27 during movement; p<0.05), independently of the chosen re-referencing configuration (Figure S3A). The early component did not show any significant modulation (Figure S3B).

To dig deeper into the circuit mechanism underlying the responses evoked by cortical stimulation, we switched to laboratory mice. Following the procedure described in Claar, Rembado et al.^21^, we recorded global neural signals with an EEG array simultaneously with up to three Neuropixels 1.0 probes^38^ in awake head-fixed mice on a freely moving wheel. Close to the Neuropixels probe in secondary motor cortex (MOs), a bipolar wire electrode was intra-cortically inserted, targeting the deep layers of MOs (layer 5/6: 1.06±0.05 mm below the surface). It repeatedly delivered single pulse ES while we measured the electrophysiological responses evoked across different locations and scales (Figure 1C, D, E). Pulses were delivered at three different stimulation intensities: low (minimum intensity to elicit a visible response at single-trial level), high (maximum intensity <100 mA not eliciting movements), medium (the average between low and high intensity). One Neuropixels probe was placed in the sensorimotor-related thalamic nuclei (SM-TH; see full list of thalamic nuclei in the Methods). The stimulation of deep layers in MOs evoked robust responses in EEG, LFP, and current source density (CSD), accompanied by a stereotyped triphasic spiking pattern characterized by an initial excitation (within 10 ms), followed by an off period (i.e., neuronal silence; 10-140 ms) and a strong rebound excitation both in MOs and SM-TH (140-250 ms; Figure 2A)^21^. As mice were free to run or to remain stationary, we classified trials based on wheel speed as either movement or rest trials (wheel speed greater or smaller than 0.1 cm/sec, respectively) and computed the average evoked potentials for EEG, LFP, and CSD for both conditions. We evaluated the early component (3- 50 ms from stimulation onset) and the late component peaking between 150-250 ms from stimulation onset and coinciding with the rebound excitation^21^. The resting condition showed significantly larger amplitude and PLF in the late time window for both global (EEG) and local (LFP and CSD) signals when high and medium current intensities were applied compared to movement (Figures 1C, D, E, S4). The modulation was reduced, or not observed, when low current intensity was applied (Figure S4). CSD analysis in MOs showed a current sink between layers 2/3 and 5 at the same time as the late response for high and medium, but not low current intensity (Figures 1E, S4C). The sink observed in this location suggests that the late component may reflect a thalamo-cortical input, given the connectivity profile of thalamo-cortical neurons^39,40^. The modulation was specific for the late EP component as opposed to the early component (Figure S5), and network specific (Figure S6).

**Figure 2.**
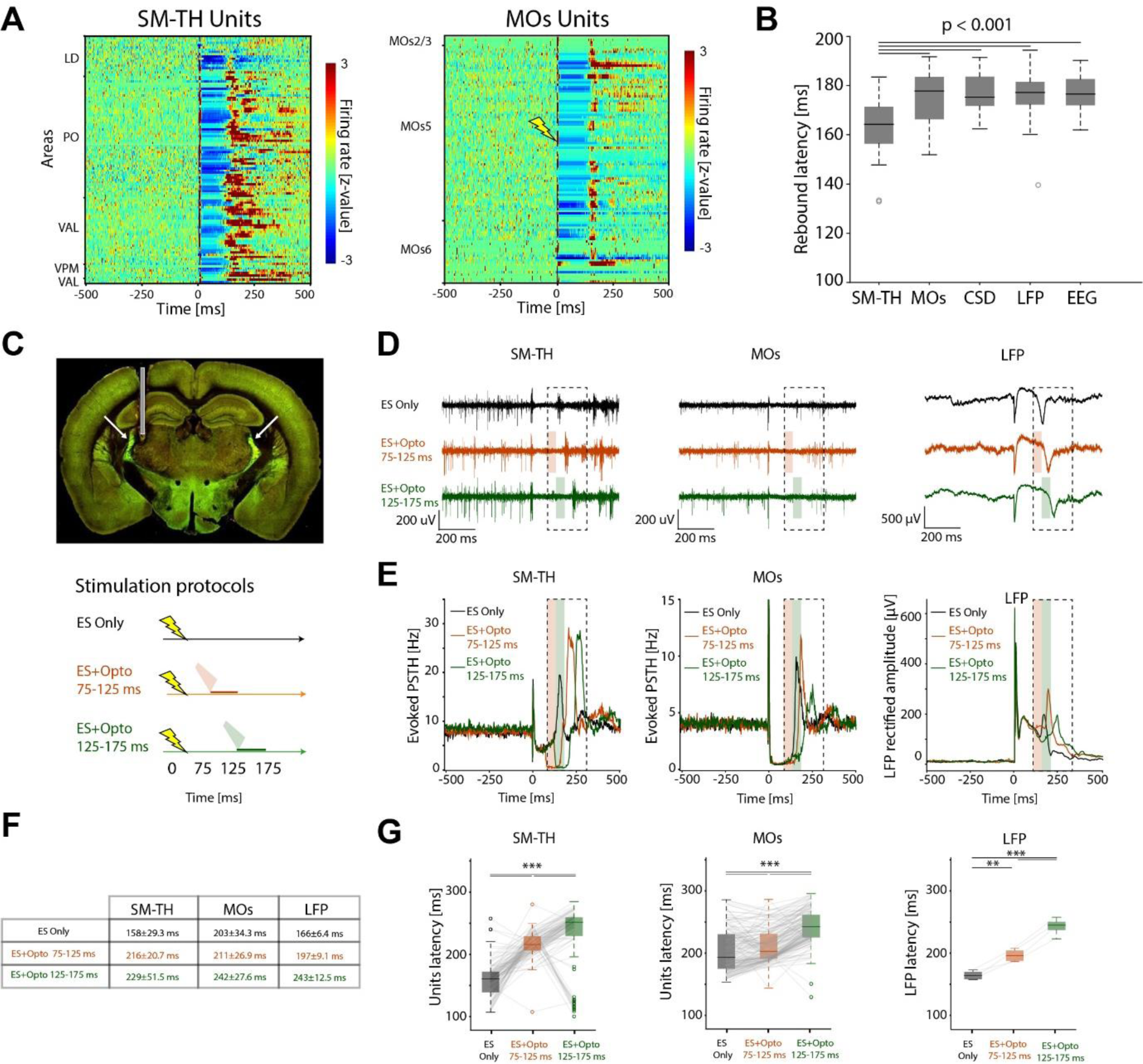
Optogenetic-induced thalamic inhibition interferes with the late cortical response to electrical stimulation of MOs. A) Responses evoked by deep MOs electrical stimulation (lightning symbol) in single units recorded by the Neuropixels probes in SM-TH (left) and MOs (right). Normalized firing rate, reported as a z-score of the average, pre-stimulus firing rate, of all neurons (only relay neurons in SM-TH and only RS neurons in MOs) recorded by the Neuropixels probes targeting the area of interest. (LD = lateral dorsal thalamic nucleus; PO = posterior thalamic nucleus; VAL = ventroanterior lateral thalamic nucleus; VPM = ventroposterior medial thalamic nucleus). B) Timing of the late responses during rest across different electrophysiological signals to MOs electrical stimulation. Global mean latencies of SM-TH and MOs units (calculated for each subject as the median across units of the first spike in the late response window and then averaged across subjects) and latencies of the CSD, LFP, and EEG peaks (two-way ANOVA. For separation between rest and movement, see Figure S7A; Tukey post-hoc test; p<0.001 for all significance lines shown in the figure; p>0.05 or n.s. for all the other comparisons). C) Top: histology of one representative VGAT-ChR2-YFP/wt mouse implanted with an optic fiber (grey rectangle). The green staining shows the intense expression of ChR2 in GABAergic neurons, particularly in the TRN (white arrows). Bottom: implemented protocols which combine electrical and timed optogenetic stimulation to causally demonstrate the thalamic origin of the late response. Three different protocols were tested: MOs electrical stimulation (ES) alone (first row, black); ES followed by 50 ms optogenetic excitation of GABAergic neurons at 75 ms (middle row, orange) and at 125 ms (bottom row, green) after electrical pulse. D) Representative raw traces of the single-trial responses evoked by the stimulation protocols shown in C (same color code) for Neuropixels electrodes in SM-TH and MOs (action potential band) and LFP in MOs (from left to right). The black dashed boxes indicate the response window used to evaluate the latency of the first spike and LFP peak (75-300 ms). E) Peristimulus time histogram (PSTH) for 118 SM-TH (left) and 155 MOs (middle) units and MOs-LFP responses averaged across 4 mice. Color code as in C and D. Note: the optogenetic stimulation specifically increased the firing rate of putative TRN units which reduced thalamic firing from 4.10 Hz to 0.25 Hz within the first 10 ms following light onset in both optogenetic conditions (Figure S20). F) Latency values (mean ± standard deviation) for SM-TH units, MOs units, and MOs-LFP peak responses for the stimulation protocols shown C. The same quantifications separated by rest and movement are reported in Table S3. G) Units’ latencies of the thalamic and cortical responses and peak latencies of MOs-LFP responses plotted in E and elicited by the stimulation protocols shown in C (Units: Wilcoxon signed rank tests; p < 0.01; LFP: one-way ANOVA test p<0.001, t-test post-hoc comparisons).

### Optogenetic dissection of the circuit causally proved that the thalamus is necessary to originate the late response to MOs stimulation in mice

Deep MOs electrical stimulation elicited a stereotyped triphasic spiking pattern – a brief excitation followed by 127.8±4.1 ms of silence and a rebound excitation – in local cortical and thalamic neurons^21^ (Figures 2A, S10B). We classified cortical units according to their spike width as regular spiking (RS) and fast spiking (FS) units (putative excitatory and inhibitory neurons, respectively; see Methods for details). Similarly, we classified thalamic units as putative SM-TH relay and reticular thalamic (TRN) units. For both resting and moving, SM-TH units rebound consistently and significantly preceded RS cortical rebound spiking and the global mean latencies of CSD, LFP and EEG peaks (p<0.001, Figures 2B, S7A). All these metrics positively and significantly correlated with the latency of SM-TH units (Figure S7B; Mixed effect models, all p<0.001). This observation points to a critical role of the thalamus in initiating the late response and generating the late EP components. To causally demonstrate such a link, we combined cortical electrical stimulation with precisely timed optogenetic inhibition of the thalamus (Figure 2C) in transgenic mice expressing Channelrhodopsin-2 (ChR2)^41^ in GABAergic neurons (VGAT-ChR2-YFP/wt). These animals were implanted with a fiber optic cannula above the TRN, considered to be the major GABAergic input to thalamic relay neurons^42–45^ (Figure 2C). Accordingly, optogenetic excitation of TRN generated a controlled suppression of the SM-TH units, inhibiting SM-TH at a precise time during the response to ES. Notably, the optogenetic stimulation activated putative TRN units which reduced the spontaneous activity of SM-TH and MOs to almost zero within 10 ms (Figs. S8A, S8C).

Compared to electrical stimulation alone, the optogenetic inhibition of SM-TH between 75-125 and 125-175 ms after the electrical stimulus progressively delayed both thalamic and cortical units’ responses (Figure 2C, D). Specifically, thalamic inhibition consistently and progressively delayed the first rebound spike (defined as the first spike after the off-period, in the 75-250 ms time window) at thalamic and cortical levels, as well as the late LFP peak (Figure 2E, F, G; Units: Wilcoxon signed rank tests, p < 0.01; LFP: one-way ANOVA test p<0.001, t-test post-hoc comparisons). On average, optogenetic stimulation between 75-125 ms following the electrical stimulation delayed the response of SM-TH units, MOs units, and the peak of the LFP by 58±39, 8±28, and 31±10 ms, respectively. When thalamic inhibition occurred 50 ms later,between 125-175 ms following electrical stimulation, response of SM-TH units, MOs units, and LFP peak were delayed by 72±53, 39±33, and 77±14 ms, respectively.

Tellingly, for all tested protocols, the firing of thalamic units in the late window always preceded activity of cortical units (Figure 2G). On the other hand, optogenetic stimulation of GABAergic neurons in MOs minimally affected the rebound response (Figure S8B; MOs: ES Only=208±39 ms; ES+Opto75-125=213±40 ms; ES+Opto125-175=220±44 ms; SM-TH: ES Only=201±36 ms; ES+Opto75-125=191±32 ms; ES+Opto125-175=204±36 ms). However, optogenetic activation alone of GABAergic units in MOs did induce a suppression of firing rate in both RS MOs and SM-TH units (Figure S8C).

Overall, these causal manipulations demonstrated that the thalamus is necessary to originate the rebound activity following electrical stimulation in mice.

### EP late component to MOs stimulation correlates with thalamic state-dependent unit synchronization

Upon further investigating rebound activity of thalamic units, we found that the evoked firing rate in MOs and SM-TH units was not modulated by movement, unlike EEG, LFP, and CSD responses. Therefore, MOs and SM-TH firing rate changes did not explain the movement-related modulation of the EPs (Figures, 3A, S9, S10 p=0.03, rest lower than movement). Instead, we found that the rectified amplitude of the EP late component was correlated to the synchronicity and bursting frequency of thalamic units (see below for more details, Figure S13). These factors were modulated by their baseline firing rate, which in turn was modulated by movement (Figures 3, S13). This observation became apparent after clustering the evoked inter-spike-interval (evoked ISI) in the late response time window of thalamic relay units (Figure 3B, C). Given the clear bimodal distribution of the evoked ISI response of relay neurons (Figure 3C), as opposed to a unimodal distribution of the cortical RS, cortical FS, and TRN units (Figure S11A; Dip test; p<0.001 for putative thalamic relay units; p>0.05, ns for all the other groups), the data-driven cluster analysis (Elbow method, Calinski-Harabasz score, and Davies-Bouldin score; metrics.calinski_harabasz_score and metrics.davies_bouldin_score functions from sklearn - Python) identified two identical thalamic cell-types (Figure S11) that responded with two distinct types of rebound activity. One with a short- evoked ISI (< 17.88 ms), referred to as high-firing units (HF; n=569), and a second one characterized by a long-evoked ISI (> 17.88 ms) and referred to as low-firing units (LF; n=700) (Figure 3C, D, E). HF units responded earlier and with less variability than LF units (Figure S11C) during the early phase (3- 50 ms), thus suggesting that HF units may receive more effective input from the cortex.

**Figure 3.**
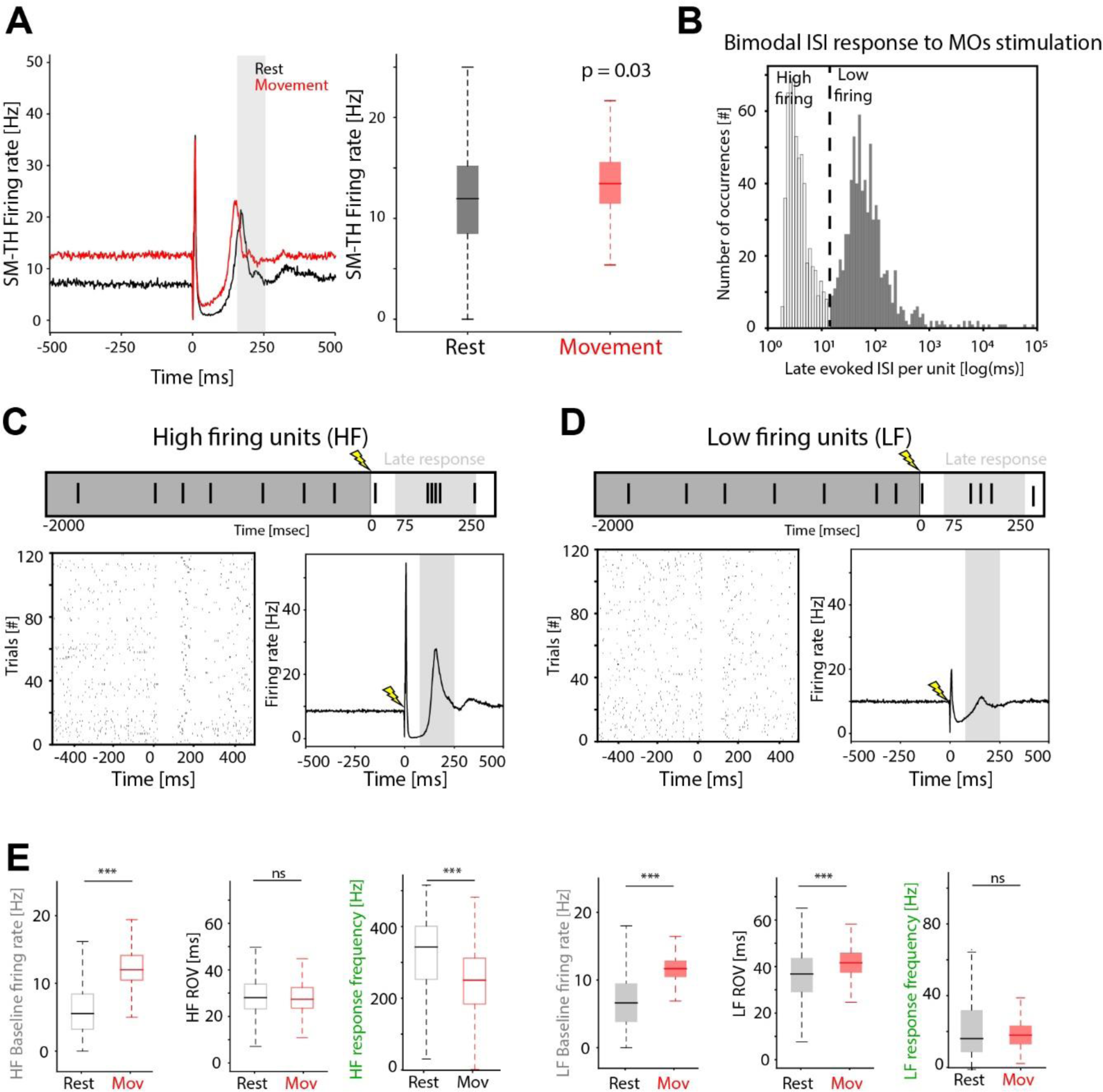
Behavioral-state-dependent modulation of rebound thalamic units’ synchronization evoked by MOs electrical stimulation underlies the modulation of the late EP component in mice. A) Modulation of SM-TH units’ evoked firing rate. Left: Evoked SM-TH firing rate at the population level during rest and movement (black and red, respectively;1269 units from 21 subjects). Right: SM-TH firing rate averaged over the late response (150-250 ms; shaded gray) at rest and movement across subjects (Wilcoxon signed rank test p = 0.03; movement larger than rest). B) Inter spike interval (ISI) histogram of the late responses (150-250 ms) evoked by MOs electrical stimulation in thalamic putative relay units (1269 units from 21 subjects). The bimodal distribution identifies two distinct response patterns: high-firing (HF, grey empty bars; 569 units) and low-firing (LF, grey filled bars; 700 units) neurons. The vertical line represents the threshold used to separate the two response patterns centered at 17.88 ms. C) Response dynamic of HF units. Top: Schematic representation of a single trial raster plot. Bottom-left: raster plot of evoked responses by electrical stimulation of a representative HF unit. Bottom-right: grand-average evoked response for all 569 HF units (21 subjects) peaking within the late response time window (150-250 ms, shaded area). D) Response dynamic of LF units (LF). Plots are structured in the same fashion as C (700 units, 21 subjects). E) Putative thalamic relay units’ features comparisons between rest (black) and movement (red) across subjects. Left: Average baseline firing rate, evoked ROV and evoked response frequency at rest and movement across HF units (Wilcoxon signed rank test, Bonferroni corrected; Baseline firing rate: p<0.001; Evoked ROV: p=0.37, ns; Response frequency: p<0.001; empty boxes). Right: the same quantifications as D visualized with filled boxplots are shown for LF units (Wilcoxon signed rank tests; Baseline firing rate: p<0.001; Evoked response variability: p<0.001; Evoked ROV: p=0.08, ns; filled boxes).

Furthermore, the strength of the electrical stimulus (i.e., current intensity) affected not only the overall number of responsive units, but also the relative size of the two subsets, such that higher stimulation intensities showed progressively more HF units than LF units (Figure S12), confirming that these two subsets reflected two different response patterns to the cortical stimulation rather than two different cell types.

The late excitatory response was characterized in terms of synchronicity, quantified for both HF and LF units as response onset variability (ROV), calculated at the single-trial level as the standard deviation of the rebound onset latency across units; and response frequency, calculated as the inverse of the evoked ISI within units (Figure S13A). By correlating baseline firing rate, ROV, response frequency and late EP component amplitude at the EEG level, we found that movement increased the thalamic baseline firing rate (from 6.78±4.4 Hz at rest to 12.48±4.6 Hz during movement; p<0.001), leading to an increased onset variability of LF units (from 35.7±11.8 ms at rest to 40.7±8.2 ms during movement; p<0.001) and to a reduced response frequency of HF units (from 309±126 Hz at rest to 241±116 Hz during movement; p<0.001). Together, these two factors reduced the amplitude of the late EP component during movement compared to rest (Figures 1, 3, S13).

### Cellular mechanisms underlying the stereotyped pause-rebound response to cortical electrical stimulation

To better understand the mechanism underlying the responses evoked by electrical stimulation, we used a computational model of the thalamo-cortical system previously developed ^46–48^ (Figure 4A). This network model is Hodgkin-Huxley type; it simulates realistic thalamo-cortical interactions and includes a rich set of parameters, including cortico-cortical, cortico-thalamic, and thalamo-cortical synaptic weights, as well as conductances for different voltage and Ca^2+^ gated currents. The model allows us to simulate rest and movement conditions by changing the input and neuromodulation^49–51^. Specifically, we simulated the movement condition by increasing the excitability of all thalamo- cortical (**TC**; corresponding to the *in vivo* SM-TH units) neurons, decreasing K^+^ leak currents and increasing the cortico-thalamic^52^ and cortico-reticular stochastic inputs simulated as Poisson process administered in AMPA synapses connecting cortico-thalamic (**PY**, corresponding to the *in vivo* RS cortical units) neurons to TC and TRN neurons^53^, compared to the rest condition (15 trials per condition).

**Figure 4.**
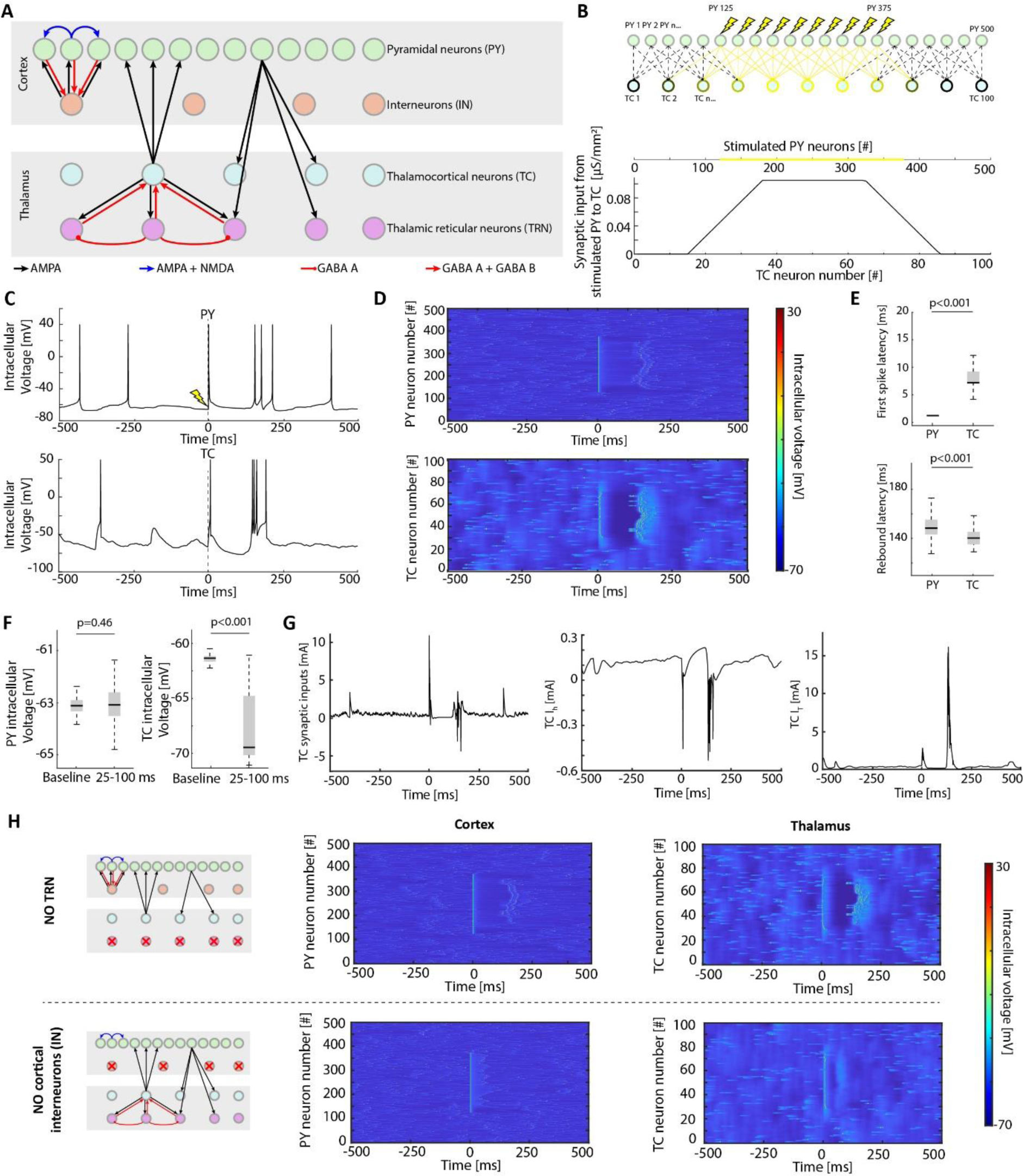
Biophysically realistic simulation of the response to cortical electrical stimulation recapitulates the *in vivo* results and provides potential explanatory mechanisms underlying the stereotyped evoked activity. A) Schematic representation of the structure of the biophysically realistic model used to generate the simulation, modified by Krishnan et al. (Krishnan, eLife, 2016). B) Schematic representation of cortical electrical stimulation and its effects on simulated TC neurons. The electrical stimulation (lightning bolts), applied to the centered half of cortical neurons (both PY and IN), was simulated as an impulsive current administered to the neuronal membrane (duration: 1 ms; intensity: 100 uA). The TC neurons in the center receive cortical inputs from several directly connected stimulated PY neurons while the TC neurons at the periphery of the stimulated area receive fewer cortical inputs (top). This is reflected in terms of cumulative synaptic inputs of the TC neurons from the stimulated PY neurons (bottom). C) Representative intracellular voltage for one PY neuron (top) and one TC neuron (bottom) as a function of time from the stimulation onset. D) Intracellular voltage for one representative trial for all PY (top) and TC (bottom) neurons. E) Median latency of the first evoked spike in the early response (0-50 ms from stimulation onset, top) and in the rebound response (75-250 ms from stimulation onset, bottom) for PY and TC neurons (early responses: PY: 3.15±3.22 ms; TC: 8.38±7.03 ms; Wilcoxon rank sum test, p<0.001. Rebound responses: PY: 151.56±11.83 ms; TC: 141.80±9.68 ms; Wilcoxon rank sum test, p<0.001). F) Median intracellular voltage during baseline and during 25-100 ms response window for PY neurons (top) and TC neurons (bottom). The intracellular voltage of PY neurons is not affected in the response window compared to baseline (64 trials, constituted by 32 rest trials and 32 movement trials; Baseline: -63.08±0.32 mV; Response: -63.16±0.88 mV; Wilcoxon rank sum test, p=0.46), while TC neurons show a significant decrease of 6.26 mV (Baseline: - 61.38±0.36 mV; Response: -67.64±3.13 mV; Wilcoxon rank sum test, p<0.001). G) Intrinsic current dynamics of TC neurons over time from the stimulation onset (0 ms). Left to right: total synaptic inputs, h-current, and T-current for a representative TC neuron as a function of time from stimulation onset. H) Simulated responses evoked by cortical stimulation in absence of either TRN or cortical GABAergic neurons. Top: schematic representation of the simulation when TRN neurons were silenced and the associated intracellular voltage for one representative trial for cortical (left) and thalamic (right) neurons. Note the preservation of the evoked stereotyped triphasic response in cortex and thalamus. Bottom: schematic representation of the simulation when GABAergic cortical neurons were silenced and the associated intracellular voltage for one representative trial for PY (left) and TC (right) neurons. In this case the evoked responses no longer show the triphasic pattern, although the thalamus still shows a period of hyperpolarization probably induced by an inhibitory effect of TRN directly activated by the electrical stimulus.

To replicate the ES protocol, we imposed a depolarizing current of 100 mA for 1 ms to half of the cortical neurons (250 PY and 50 interneurons (**IN**) spatially located in the center of the network (Figure 4B). Each TC neuron received cortical inputs from the closest PY neurons within a certain radius (Figure 4B). So, the TC neurons in the center of the network received inputs from several stimulated PY neurons with respect to the TC neurons at the periphery of the simulation (Figure 4B). This was reflected in terms of cumulative synaptic inputs from the stimulated PY neurons to TC neurons, showing that the stimulated PY neurons exert maximum effect on thalamic neurons in the center of the network, and progressively decreased towards the borders of the stimulated area (Figure 4B).

The model successfully recapitulated the triphasic response characteristics observed in the mouse (Figures 3A;^21^, 4D, S9A). Specifically, it also showed that during the cortical and thalamic off-periods, between 25 and 100 ms after stimulation, there was no significant change, compared to baseline, in the membrane potential of PY neurons (p=0.46; Baseline: -63.08±0.32 mV; Response: -63.16±0.88 mV, Figure 4C, 4F left) but the membrane potentials of the TC neurons were significantly hyperpolarized (p<0.001; Baseline: -61.38±0.36 mV; Response: -67.64±3.13 mV; Figure 4C, 4F right).

This hyperpolarization, which followed an early spike of the same cells due to the stimulation driven cortical input, was caused jointly by a reduction of cortico-thalamic excitatory inputs and inhibition from the TRN cells (Figure 4G, left). As the hyperpolarization gradually increased (reaching values lower than -75mV), it was accompanied by (a) de-inactivation of the low-threshold Ca2+ (T-) currents and (b) slow activation of the h-currents (Figure 4G, center). The activation of the h-currents then led to the depolarization and quick activation of the T-currents triggering low-threshold Ca2+ spike (Figure 4G, right), as previously described^54^. To further clarify the mechanisms allowing thalamic hyperpolarization, we performed additional simulations in absence of TRN and cortical inhibitory neurons (Figure 4H). When TRN was silenced, the pause-burst pattern was largely unaffected (Figure 4H, top). Conversely, when cortical interneurons were silenced, PY showed a reduced off-period, and the thalamic rebound was abolished. However, the thalamus did show a period of hyperpolarization probably induced by TRN (Figure 4H, bottom). Our results suggested that the GABA-mediated cortical off period is critical for the occurrence of the thalamic rebound response although TRN may also contribute to it; the reduced excitatory inputs to the thalamus lead to transient hyperpolarization of the thalamic TC neurons, which then allows for the generation of the thalamic rebound. Importantly, our optogenetic experiments confirmed that the activation of cortical GABAergic units alone is sufficient to induce both cortical and thalamic off-periods (Figure S8C).

Lastly, as suggested by the mouse data, the model confirms that TC neurons exhibit two rebound regimens, determined by the amount of synaptic inputs that they receive from cortical neurons (Wilcoxon ranksum test; p<0.001; HF: 0.77±0.098 μS/cm^2^; LF: 0.37±0.16 μS/cm^2^; Figure S14) and that the differences in the thalamic rebound during rest and movement depends on the hyperpolarization level of TC neurons (Figure S14). Indeed, the movement condition led to a relative depolarization of TC neurons, so that the reduction of cortical input was insufficient to fully de-inactivate T-channels in TC neurons (Figure S14).

## Discussion

We demonstrate that the thalamus is a major contributor to the late EP component elicited by cortical stimulation in mice and provide compelling evidence that denotes its role in shaping very similar responses in humans. In mice, we proved that the thalamus is necessary to generate this late component via a causal manipulation of the cortico-thalamic circuit by combining electrical stimulation with optogenetic inhibition (Figure 2). Given the striking similarities in the EP waveforms (Figures 1, 5A) and the comparable behaviorally modulated late components in humans, we infer that similar mechanisms underly the EPs in both species. Specifically, the state of the subject modulates the degree of spike synchronicity and burst frequency of the thalamic evoked rebound responses, which in turn modulate the amplitude of the late EP component captured by the EEG electrodes (Figure 3). A biophysical simulation of the cortico-thalamic circuit^46^ predicted that these stereotyped evoked responses are mediated by the intrinsic dynamics of thalamic currents in conjunction with thalamic and cortical GABAergic neurons (Figure 4).

### The key role of the thalamus in generating the late EP

Electrical stimulation of the deep layers of MOs in mice elicits a stereotyped triphasic spiking pattern in local cortical and thalamic neurons that resembles the responses evoked at the EEG level^21^. We found that the late EP component, between 150 and 250 ms from stimulation onset, is modulated by the behavioral state, such that its magnitude decreases in running animals, not only at the EEG level, but also at the LFP and CSD levels (Figure 1). We used timed optogenetic manipulations of the circuit to prove that this component depends on thalamic activity (Figure 2), but not cortical activity (Figure S8). Like the late EP component (Figure 1), the pause-burst pattern in the thalamus is state-dependent: while mice are running, the thalamus is more depolarized (high baseline firing rate, Figures 1, 3) compared to resting. As shown using a biophysical network model^46^, running reduces stimulation-evoked hyperpolarization in thalamo-cortical neurons (Figure S14) that is necessary to de-inactivate the voltage-gated, T-current^55^, and evoke the subsequent spike bursts^56–59^. These currents decrease the bursting frequency and population synchronicity of the rebound compared to rest (Figures 3, S14). Previous work showed that the state-dependent transition of TC neurons^58^ from burst firing to single spiking mode, is mediated by metabotropic glutamate receptors^60^. These two neuronal patterns of activity are often associated with sleep and arousal, respectively^61^, although bursting may sporadically occur during wakefulness^62,63^. Our results show that the animal’s state (Figures 1, 3) and the intensity of the ES (Figure S4) modulate the late EP component via their effects on thalamic bursting frequency and synchronicity, rather than cortical and thalamic evoked firing rate (Figure 3, S9).

A similar state-dependent modulation is observed in humans, such that when the stimulated cortico- thalamic network is activated by movement, either active or passive, the late EP component measured by EEG (or iEEG) is smaller than at rest (Figures 1, S2, S3). The modulation observed during passive movement suggests that peripheral inputs are sufficient to change the state of the thalamus. Importantly, like in mice (Figure S6), the modulation of the rebound in humans is region specific, as it is not observed if the stimulation is delivered to a posterior cortical area not directly engaged by the task (Figure S2), and is specific to the late, rather than the early, EP component (Figures S2, S3, S5), emphasizing their different mechanistic origins.

There are many differences between homo sapiens and mus musculus, including neuronal population, gene expression, and brain size^64–66^. The difference in brain size should affect the timing of signal conduction from thalamus to cortex and *vice versa*. However, given an estimated conduction velocity of 5-50 m/s^67^, the difference in thalamo-cortical conduction times between the two species will be below 10 ms, significantly smaller than the 100 ms time window that defines the late EP component in this study. Compatible with our results, however, are the several similarities across these two mammalian species, ranging from common thalamic burst dynamics^68–71^ and remarkable conservation of brain rhythms^72^ to shared features of thalamocortical and corticocortical connections^73^. Moreover, we focused on the presence of the state-dependent modulation per se, rather than comparing the absolute magnitude of the modulation, expected to be different due to the differences in recording and stimulating modalities in mice and humans. We defined movement for mice and humans in different ways: running and actively squeezing a rubber ball or passively moving the same hand, respectively. To account for the different definition of movement, for the experiments in humans, we stimulated the premotor cortex contralateral to the hand engaged by the active or passive task, thus realistically comparing an active condition of the perturbed cortico- thalamic network to a resting state.

### Mechanisms of neural responses to cortical stimulation

We previously showed^21^ that deep ES stimulates pyramidal cells, including thalamic projecting ones. This generates an initial excitation, both in cortex and thalamus, followed by an off-period and a rebound excitation initiated by the thalamus^21^ (Figure 2). This rebound is characterized by bursting dynamics (Figure 3), mediated by the intrinsic dynamic of T-currents, in thalamic relay cells^55^. T-currents are activated by the depolarizing h-current, primarily mediated by HCN channels^74–76^, which in turn is activated by pronounced hyperpolarization^24,63,77–80^ (Figure 4G). According to our model, the cortical stimulation significantly hyperpolarizes the thalamus – as opposed to the cortex, whose intracellular voltage is comparable to baseline (Figure 4F) – by both withdrawal of excitation due to the cortical silence and disynaptic inhibition via stimulation of TRN neurons, the primary source of GABAergic inputs to the relay thalamic nuclei^43–45,74^ (Figure 5B). Similarly, the cortical off-period may be due to a transient inhibition mediated by GABAergic neurons, and/or a lack of excitatory input from the thalamus. Because cortical and thalamic areas sustain each other’s firing through recurrent loops (Figure 5B), it is difficult to isolate the individual effects. Indeed, a suppression of either cortex or thalamus by directly activating their GABAergic neurons induced a decrease of firing rate in both areas (Figure S8C).

**Figure 5.**
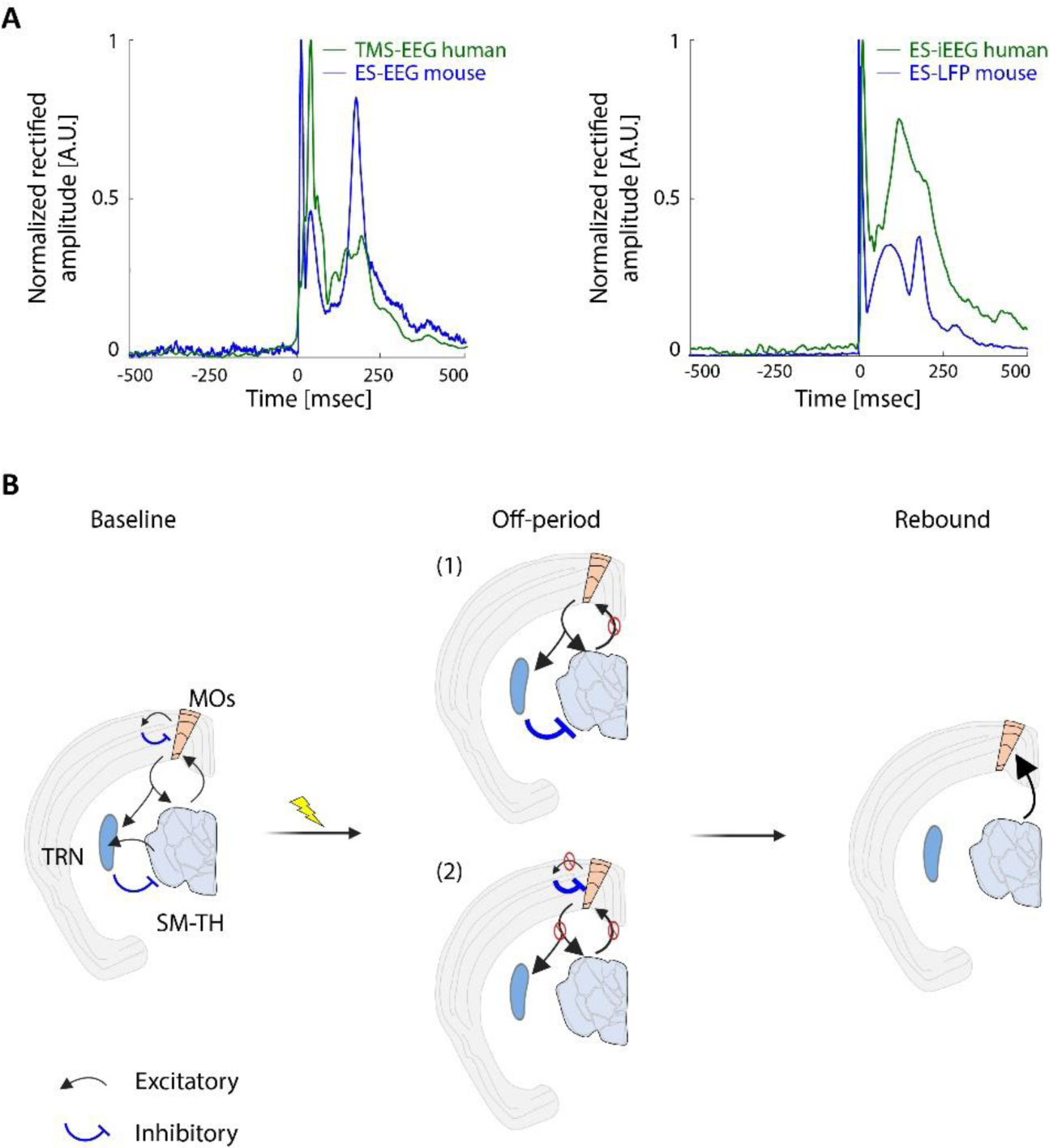
Similar mechanisms in mouse and human explain the evoked responses following invasive and non-invasive stimulation. A) Evoked responses elicited by cortical stimulation (invasive and non-invasive) are remarkably similar between human and mouse. Left: grand average EEG evoked responses rectified and normalized by their maximum value elicited by TMS in humans (green, 12 subjects) and by ES in mice (blue, 15 subjects). Right: grand average iEEG and LFP evoked responses rectified and normalized by their maximum value elicited by ES in humans (green, 5 subjects) and mice (blue, 13 subjects). B) Diagram of the thalamocortical circuits at baseline characterized by recurrent inhibitory and excitatory loops within secondary motor cortex (MOs), between MOs and somatomotor thalamus (SM-TH) and thalamic reticular nucleus (TRN). Possible mechanisms underlying the off-period following cortical stimulation. In scenario (1), the cortical stimulation activates TRN which in turn suppresses SM-TH via a volley of GABAergic inputs which causes the thalamic off-period and in turn the cortical off-period. In scenario (2), the early excitatory response in the cortex induces a GABAergic-mediated cortical off-period, which interrupts the recurrent loops sustaining the activity in cortex and thalamus, ultimately inducing thalamic and cortical off-periods. The two scenarios are not mutually exclusive and could synergistically occur as shown by our simulated and *in vivo* data. In both cases, the thalamic hyperpolarization engages low-threshold calcium currents, It, which initiate the thalamic rebound response and the consequent cortical rebound.

As discussed previously, cortical stimulation generates an initial excitation locally (including in GABAergic neurons) and in the thalamus, followed by an off-period in both areas. When we ran the simulation in the absence of cortical GABAergic neurons, we did not observe the evoked pause-rebound pattern in either cortex or thalamus (Figure 4H, bottom), suggesting that the cortical GABAergic neurons are critical for generating the stimulation-induced cortical and thalamic off-period. Their key role in deactivating recurrent local activity after an electrical stimulus has been shown before in cortical slices^81^. This induced local down state could trigger a down state in the directly connected thalamic nuclei by reducing the cortical inputs to them, hence causing an off-period in the thalamus. In line with previous findings^82^, cortical stimulation directly activates a subset of TRN neurons, as supported by TRN recordings from six animals where only two responded to the stimulation, and they responded with a stereotyped triphasic pattern (Figure S10B). Our model shows that TRN neurons are not necessary to evoke the thalamic off-period (Figure 4H, top). Though the thalamus still hyperpolarized following ES when the cortical GABAergic neurons were silenced (Figure 4H, bottom) possibly due to direct engagement of TRN, confirming its contribution to the thalamic off-period. Overall, our results suggest that GABAergic neurons in both cortex and thalamus concurrently contribute to generate the thalamic off-period.

Following the off-period, we observe a rebound response in the thalamus, characterized by spike bursts, and a subsequent rebound in the cortex. We showed that when TRN is optogenetically activated (Figures 2, S8), it interferes with the electrically evoked pause-burst dynamic by further suppressing the thalamic relay neurons (Figure S8). A possible explanation is that optogenetic TRN activation keeps thalamic relay cells hyperpolarized such that the h-current is not sufficient to depolarize the membrane, thus delaying the rebound burst (low-threshold Ca^2+^ spike). Once the optogenetic stimulation of TRN neurons ends, the thalamic relay cells respond with an even larger rebound activation (Figure 2B). With the thalamus optically suppressed, the cortex cannot generate an independent rebound, extending the evoked off-period over 240 ms, hence delaying the rebound both in cortex and thalamus (Figure 2). On the other hand, when ES was combined with direct optogenetic activation of cortical GABAergic neurons, we did not cause a similar delay of the rebound response in either cortex or thalamus. Taken together these two observations causally demonstrate the thalamic dependency of the late rebound response to ES (Figure S8). Thus, we propose that both cortical and thalamic GABAergic neurons contribute to hyperpolarizing the thalamus after cortical stimulation, thus enabling the thalamic rebound response (Figure 5B).

Another important finding of the study is that cortical stimulation induces two different thalamic response patterns, here called high firing (HF) and low firing (LF) (Figure 3) in both *in vivo* and simulated data (Figure S14). Rather than reflecting different neuronal types (Figure S11), these two response patterns are explained by the connectivity profile of each neuron (Figure S14). Thalamic neurons receiving large inputs from the stimulated cortical area showed a HF response (Figure S14), and they responded more promptly and with less variability than thalamic neurons receiving smaller inputs from the stimulated cortical area and characterized by a LF response (Figure S11C). *In vivo* data further supported this prediction, showing that the intensity of the electrical stimulus affected not only the overall number of responsive units, but also the relative size of the two subsets, such that higher current intensities showed a progressive shift from LF to HF response patterns (Figure S12).

Intriguingly, the similarities in the stimulation-evoked responses and their behavior-dependent modulation are observed across stimulation modalities (Figures 1, 5A). In line with this observation, TMS in non-human primates evokes stereotyped triphasic spiking responses^27^ and it engages GABAergic cortical neurons^83^. The origin of the late EEG responses evoked by TMS in humans have been a source of controversy, given that multisensory peripheral factors may contribute to their generation^20^. However, our control experiments (Figure S2) showed that the observed modulation was not due to sensory responses to TMS^84^. This result confirms that the late components evoked by confounds-controlled TMS are likely generated by direct cortical activation that engages similar mechanism as the one elicited by invasive ES in humans and mice (Figures 1, 5). TMS and ES activate the underlying tissue through different mechanisms^30,85–92^, which may explain why the early component evoked by TMS (Figure S2) is modulated by movement, but the one evoked by ES in mice and humans is not (Figures S3, S5). However, early and late components in the TMS-EEG experiments are independently modulated (Figure S2), thus supporting the hypothesis that they originate from different mechanisms. We therefore conclude that both TMS and ES directly engage the thalamus.

### Implications of the study

Overall, our *in vivo* and *in silico* results (Figures 2,3,4) suggest that the main actors of the late responses to cortical stimulation (invasive and non-invasive) are the intrinsic thalamic currents, I_h_ and I_t_, in thalamo-cortical cells and the cortical and thalamic GABAergic neurons. As these are similar across species, from mice to non-human primates to humans, we surmise that the neural responses to cortical stimulation originate from highly preserved mechanisms, including the thalamic dependency of the late EP component^68–71^. Note that this does not exclude cortico-cortical contributions. Given that we demonstrated that the thalamus plays a critical role in shaping these responses, it is reasonable to expect that it also plays a major role in the effectiveness of all the research, clinical and neuroprosthetic applications based on cortical stimulation^4,7,11,12,17,18,93–97^.

The thalamic state-dependency of the late EP component (Figure 3) suggests that the thalamic feedback that reaches the cortex is integrated with information related to the state of the thalamus. We therefore propose that this late EP may be used as a non-invasive indicator of the state of the thalamus, potentially providing a new biomarker for thalamo-cortical (dys)functions.

## Acknowledgments

We sincerely thank David McCormick for the generous feedback and the extended discussions. We thank Josh Siegle for technical advice, and comments on the manuscript. We thank the clinical and surgical team of Niguarda Hospital (Milan, MI, Italy), the patients and their families; the computational resources provided by the Core Facility INDACO, which is a project of High-Performance Computing at the Università degli studi di Milano (http://www.unimi.it) and Thierry Nieus for the technical assistance. We thank Han Hou for providing the transgenic VGAT mice and Sam Gale for directions on the optogenetic experiments; the Allen Institute Animal Care, the Neurosurgery and Behavior, and the Lab Animal Services teams for mouse husbandry, care, and habituation; the Allen Institute Manufacturing and Process Engineering team for experimental hardware and software support; and the Allen Institute Transgenic Colony Management and the Imaging teams. We wish to thank the Allen Institute founder, Paul G. Allen, for his vision, encouragement, and support.

## Contributions

Conceptualization: I.R., S.R., M.M. and C.K.; Methodology: I.R., S.R., L.C., C.K., G.K., S.S., M.R., A.P., M.M.; Formal Analysis: S.R.; Investigation: I.R., S.R., L.C., L.M., G.F., F.M.Z., G.H., M.S., M.R., I.S., A.P., E.M.; Data Curation: S.R., L.C., L.M.,I.R., F.M.Z., A.P., E.M.; Writing – Original Draft: I.R., S.R., C.K.; Writing – Review & Editing: I.R., S.R, C.K., L.C., G.K., M.R., A.P., E.M., M.B., S.S., M.M., L.M., F.M.Z., I.S.; Visualization: S.R., I.R.; Supervision: I.R., C.K., M.M., M.B., S.S., M.R., I.S., A.P.; Project administration: I.R.; Funding Acquisition: C.K., M.M.

## Materials and methods

### Mouse experiment

Mouse data has been collected through the experimental procedures described in Claar, Rembado, et al^21^. A summary of these methods and details of the procedures that differ are provided below.

#### Mice

Mice were maintained in the Allen Institute animal facility and used in accordance with protocols approved by the Allen Institute’s Institutional Animal Care and Use Committee under protocols 1703, 2003 and 2212. Experiments performing electrical stimulation alone used C57BL/6J wild-type mice (n=21), while experiments performing optogenetic manipulation used VGAT-ChR2-YFP/wt mice (n=4). Male and female of both wild-type C57BL/6J mice and VGAT-ChR2-YFP/wt mice were purchased from Jackson Laboratories (JAX stock #000664) and were 9-28 weeks old at the time of *in vivo* electrophysiological recordings.

After surgery, all mice were single-housed (reverse 12-h light cycle; temperatures 20-22 °C; humidity 30-70%; ad libitum access to food and water). All experiments were performed during the dark cycle.

#### Surgical procedures and habituation

Each mouse went through the following order of procedures prior to the day of the experiment: 1) an initial sterile surgery to implant an EEG array and a titanium headframe; 2) five days of recovery time post-surgery; 3) at least three weeks of habituation to head-fixation; 4) and a second sterile surgery to perform small craniotomies to allow for insertion of the stimulating electrode and Neuropixels probes. Refer to Claar, Rembado et al^21^ for details.

The day of the first surgery, mice undergoing optogenetic stimulation of the thalamus were implanted with an optical fiber targeting the reticular nucleus of the thalamus. After removing the skin and exposing the skull, we drilled a hole (0.5 mm in diameter) and slowly inserted a syringe needle to a depth of 3250 μm, waited for 5 minutes for the tissue to stabilize and then retracted the syringe needle and inserted the optical fiber. The optical fiber was then secured to the skull with White C&B Metabond (Parkell, Inc, Edgewood, NY, USA) together with the titanium headframe.

#### Experimental procedure: EEG and Neuropixels recordings and cortical electrical stimulation

For detailed methods see the methods section in Claar, Rembado, et al^21^. In summary: the day of the experiment, the mouse was placed on the running wheel and fixed to the headframe clamp with two set screws. A thin layer of Kwik-Cast was removed to expose the craniotomies and abundant ACSF was added on top of the skull to keep the exposed brain tissue hydrated. A 3D-printed cone was then lowered to prevent the mouse’s tail from contacting the probes and a black curtain was lowered over the front of the rig, placing the mouse in complete darkness and free to run or rest at its discretion. In addition to recording from multiple Neuropixels probes, the 30 electrode EEG array was connected to a 32-channel head-stage (RHD 32ch, Intan Technologies, Los Angeles, California) controlled by an Open Ephys acquisition board^21,98^. Electrical stimulation was delivered through a custom bipolar platinum-iridium stereotrode (Microprobes for Life Science, Gaithersburg, Maryland) consisting of two parallel monopolar electrodes (50 kOhm impedance) with a vertical offset of 300 µm between the two tips. The stimulating electrode was acutely inserted using a 3-axis micromanipulator, like the Neuropixels probes, targeting secondary motor cortex (MOs), layer 5/6 (1.4±0.24 mm below the brain surface). Up to 120 biphasic, charge-balanced, cathodic-first current pulses (200 μsec per phase, 3.5-4.5 sec jittered inter-stimulus interval) were delivered at three different current intensities (up to 360 pulses total). The current intensities were chosen for each animal before starting the experiment based on the following criteria: 1) the maximum stimulation intensity was selected as the maximum intensity that did not evoke any visible twitches (below 100 μA), 2) the minimum stimulation intensity was selected as the minimum intensity that evoked a visible response for most of the EEG electrodes (n > 15) for at least 20 ms following the stimulus onset, 3) the medium stimulation intensity was selected as the average between maximum and minimum stimulation intensity (High: 66.15±12.11 μA; Intermediate: 47.35±13.30 μA; Low: 30.59±15.80 μA). For the optogenetic experiments a step function of 5 mW blue light was applied for 50 ms at 75 ms or 125 ms after the electrical stimulation onset either targeting the thalamus (Figure 2) or targeting MOs (Figure S22).

#### EEG quality control and pre-processing

Before the experiment, the EEG signals were tested by exposing the animal to visual flashes and evaluating the signal-to-noise ratio of the EEG evoked responses. Animals with low signal-to-noise ratio, high levels of 60 Hz noise, large long-lasting stimulation-related artifacts, or large movement artifacts were not included for further analyses. Experimental EEG data was then preprocessed as follows. The stimulation artifact was masked by copying the raw signal from -9 to -3 ms, reversing, and replacing it in the -3 to +3 ms artifact window. After artifact masking, EEG recordings were visually inspected to identify electrodes containing noise artifacts or remaining large and/or long-lasting stimulation artifacts. These were excluded from further analysis, removing an average of 3.9±3.7 artifact-contaminated electrodes out of 30 for each subject. EEG signals from all good electrodes were high-pass filtered (0.5 highpass 3^rd^ order Butterworth filter, signal.butter and signal.filtfilt function from Scipy – Python). Finally, the continuous EEG signals were segmented into epochs from -2 to +2 s from stimulus onset and saved for further analysis.

#### Neuropixels EPs pre-processing

Continuous Neuropixels LFP signals were segmented into epochs from -2 to +2 sec from stimulus onset, detrend (signal.detrend function from Scipy – Python), and saved for further analysis.

To compute CSD, LFP epochs underwent an automatic channel rejection based on Chebyshev’s inequality, iteratively interpolating any channel whose amplitude instantaneously exceeded ±7 standard deviations with respect to the others (similar to Russo et al^99^). The cleaned LFP voltages were smoothed in time (smoothing window=1.6 ms) and space domain (1^st^ smoothing window=26 channels; 2^nd^ smoothing window=4 channels). The CSD was calculated as the second spatial derivative^100^ from the cleaned, smoothed LFP signals. The CSD formulation employed assumes an ohmic conductive medium, constant extracellular conductivity (σ=0.3 S/m), and homogeneous in-plane neuronal activity, with the boundary condition of zero current outside the sampled area.

#### Neuropixels units pre-processing

Neuropixels AP raw signals were also artifact masked before being pre-processed and spike-sorted using Kilosort 2.0^101^ as described by Siegle, Jia et al.^38^. After spike sorting, any spikes that occurred during the artifact window (0 to +2 ms from stimulus onset) were removed from further analysis. High quality units were identified for further analysis using metrics described by Siegle, Jia et al^38^. We classified cortical regular spiking (**RS**) and fast spiking (**FS**) neurons (putative pyramidal and inhibitory neurons, respectively) based on their spike waveform duration (RS duration > 400 μs; FS duration ≤ 400 μs)^102–107^. Similarly, thalamic units were classified as putative relay neurons if their spike width was above 450 μs^108–110^ and putative reticular (TRN) neurons if their spike width was below 350 μs^110^.

### Intracerebral EEG

#### Subjects

Subjects included in the present section were selected from a population of patients affected by drug-resistant focal epilepsy undergoing presurgical screening with intracerebral EEG (iEEG) electrodes in “C. Munari” Epilepsy Surgery Center (ASST Niguarda-Ospedale Ca’ Granda, Milan, Italy). Inclusion criteria were applied as follows: (I) adult, (II) location of one or more iEEG electrodes in premotor areas, (III) epileptogenic zone located outside the stimulated premotor site, (IV) absence of neurological and psychiatric deficits preventing the execution of the tasks (for demographics and clinical details see Table S2).

All patients included in the present section provided written informed consent. The experimental protocol was approved by the local ethics committee of Milan (ID 348-24.06.2020, Milano AREA C Niguarda Hospital, Milan, Italy) in line with the Declaration of Helsinki.

#### Intracerebral EEG setup

iEEG was recorded from platinum-iridium semiflexible multi-contact intracerebral probes (diameter: 0.8 mm, contact length: 1.5mm, inter-contact-spacing 2mm; Dixi Medical, Besancon France). The number and location of the intracerebral electrodes was decided according to one or more clinical hypotheses, as described in Cossu et al and Cardinale et al^111,112^. iEEG electrodes were implanted using a robotic assistant (Neuromate, Renishaw Mayfield SA). Overall, each subject was implanted with 18±1.4 electrodes (8 to 18 contacts per electrode; 164.6±3 bipolar and 190.4±1.6 monopolar contacts per patient). iEEG signal was acquired through a 192-channels amplifier (Nihon-Kohden Neurofax-1200) and sampled at 1000 Hz. Two adjacent contacts located entirely in white matter served as reference and ground.

#### Contacts localization

For each subject, the location of intracerebral contacts was assessed by coregistering the pre-implant 3D-T1 magnetic resonance image (MRI; Achieva 1.5 T, Philips Healthcare, Holland, Amsterdam) with the post-implant Computed Tomography (CT; O-arm 1000 system, Medtronic, Ireland, Dublin) of the subject using the FLIRT software tool^113^. The MRI was processed through Freesurfer^114^, the location of each contact was estimated using the two software SEEG Assistant^115^ and 3D Slicer^116^ and assigned to an anatomical area of the Desikan-Killany atlas^117^. The anatomical location of each contact was visually confirmed by a trained neurophysiologist. Stimulated contacts in premotor areas were functionally confirmed according to their ability to induce a dystonic motor response of the contralateral hand when electrically stimulated with high-frequency stimulation (50 Hz for 5 seconds^118^.

#### Experimental procedure

Experimental procedures were performed at Niguarda Hospital under medical supervision. During the stimulation sessions, patients were sitting on a hospital bed, and they underwent video-EEG monitoring. Electrical stimulation (square positive biphasic bipolar pulses; pulse width: 0.5 ms) was administered through a Nihon-Kohden Neurofax-1200 system (Nihon Kohden, Tokyo, Japan) and delivered between two adjacent contacts pertaining to the same electrode and located in the premotor area (Broadman area 6). Each patient underwent intracerebral stimulation at rest and during self-paced intermittent hand squeezing task contralateral to the stimulated hemisphere. The order of the conditions was randomized for each patient. Depending on the clinical timeline, each stimulation session included 26.4±3.1 pulses with a stimulation frequency of 0.5 Hz. The stimulation intensity administered for each patient was selected as the highest intensity (maximum 5 mA) without inducing twitches nor subjective perceptions. No seizures occurred during the stimulation protocol.

#### Intracerebral EEG pre-processing

The stimulation artifact was removed through a tukey-filter^15^. The raw signal was filtered (1 Hz zero-phase 3^rd^ order Butterworth highpass filter; butter and filtfilt functions from Matlab) and epoched from -700 to 1300 ms around the pulse. iEEG signal was bipolar referenced by subtracting the activity of each channel to the contiguous channel. For both monopolar and bipolar recordings, all channels and trials signals were visually inspected and rejected if electrical artifacts and/or epileptic events were present. Only the signals recorded from responding contacts (exceeding 6 standard deviations of the baseline^119^) located within 3 cm from the stimulated area were retained for further analyses in the region of interest (ROI).

### TMS-EEG experiments

#### Subjects

Subjects were recruited from a population of healthy adults and screened for eligibility (for demographics see Table S1). Participants were excluded if the medical screening indicated any of the following conditions: suspected pregnancy, presence of metal parts, clinical history suggestive of epilepsy or brain lesions, cognitive impairment preventing the execution of the tasks, recent intake of drugs with neurological effects.

All subjects provided written informed consent. The experimental protocol was approved by the local ethics committee of Milan (Ethics Committee Milano Area A, Milan, Italy) in line with the Declaration of Helsinki.

#### TMS-EEG setup

TMS was performed using a 50/70 mm Air Cooled Focal figure-of-eight coil (Aircooled focal coil, Nexstim Plc, Finland) driven by a stimulator unit (NBS9 stimulator unit, Nexstim Plc, Finland). TMS pulses were triggered from an external trigger box (BrainProducts, Munich, DE) and delivered with a randomized inter-pulse-interval jittered between 2 and 2.3 seconds. The targeted site was continuously monitored on the subject 3D-T1 MRI through the real-time neuronavigation system integrated in the NBS9 system (Nexstim Plc, Helsinky, FI).

EEG signals were recorded by using TMS-compatible 64 channels EEG amplifiers (either BrainAmp MR+ and BrainAmp DC, Brain Products, Gilching, Germany) connected to a high-density 64-channel cap (EasyCap, Wörthsee, DE) whose EEG c-shaped electrodes were positioned in the standard 10-20 locations. The impedance of all electrodes was kept below 10kΩ by applying Electro-Gel (Electro-cap International, Inc., OH, US) and EEG signals were sampled at 5000 Hz. EEG reference and ground electrodes were positioned on the subjects’ forehead.

#### Experimental Procedure

Experimental procedures were performed at the University of Milan, Italy. Subjects laid on an electronically adjustable Nexstim chair (NBS9 chair, Nexstim Plc, Finland) wearing a high-density 64 channels EEG cap and in-ear earphones (HA-FX8, JVCKenwood Corporation, Yokohama, JP) for the administration of subject-specific masking noise to prevent TMS sound perception^120^. TMS was administered on the left premotor cortex (Brodmann area 6) with an inter-stimulus-interval ranging from 2 to 2.3 sec. The precise stimulation site, angle and intensity for each subject was chosen according to the criteria proposed by Casarotto et al^121^ to minimize muscular artifacts and maximize cortical responses. Once the stimulation parameters have been defined, each subject underwent 3 TMS-EEG sessions for three different conditions of the right hand (contralateral to the stimulated hemisphere): resting (rest), self-paced intermittent hand squeezing of a rubber ball (active movement), and passive movement of the same hand by an experimenter (passive movement). The order of the conditions was randomized across subjects. In a subset of 5 subjects, we acquired 6 control sessions targeting either occipital and parietal areas (Broadman areas 7,17,18, and 19). For a subset of 7 subjects, we acquired an extra premotor TMS session at rest preceding the 3 randomized sessions.

#### Median nerve stimulation

To test whether the movement-related modulation was specific to TMS evoked responses and not caused by a sensory stimulation induced by the TMS pulses themselves, we performed control experiments (10 subjects) in which we recorded the EEG response to the somatosensory stimulation of the left median nerve (somatosensory evoked potential – SSEP). The stimulation was administered through silver chloride cup electrodes placed about 5 cm apart along the median nerve on the volar side of the left forearm. The stimulated area was first scrubbed with NuPrep skin preparation gel (Weaver and Company, Denver, CO, US), then the cup electrodes were positioned using Ten20 conductive paste (Weaver and Company, Denver, CO, US) and fixed with micropore medical tape. Electrical biphasic pulses with alternating polarity were generated by an electrical stimulator (Digitimer DS10A, GB) driven by a trigger box (BrainProducts, Munich, DE) and delivered at a randomized inter-pulse-interval ranging between 2 and 2.3 sec. For each investigated condition (active movement, passive movement, and resting) we administered between 200 and 300 pulses.

The order of the conditions was randomized across subjects. Stimulation intensity was determined for each subject as 90% of the motor threshold (i.e. the threshold inducing motor twitches for 50% of the delivered pulses). For a subset of 7 subjects an extra control session at rest was collected preceding the other 3 sessions.

#### TMS-EEG and SSEP pre-processing

EEG signals pre-processing was performed using a custom-written Matlab (The MathWorks) pipeline similar to the one described by Fecchio et al^122^. For each EEG channel, we removed TMS and SSEP stimulation artifacts by replacing the signal in the 8 ms following the pulse with the mirrored signal from the 8 ms preceding the pulse. A high-pass filter was then applied (1Hz 3^rd^ order high-pass Butterworth filter, butter and filtfilt functions from Matlab). Channels and trials were visually inspected and rejected if contaminated by artifacts and the remaining good signals were re-referenced to the common average. EEG signals were then epoched from -600 to +600 ms around the pulse, and each epoch was lowpass filtered (Antialiasing Chebychev Type I IIR 8^th^ order filter) and downsampled from 5000Hz to 1250Hz (decimate function from Matlab). Independent Component Analysis (runica function from Matlab) was used to remove the following EEG artifacts: eye movements, muscle activity, TMS decay. Finally, EEG epochs were filtered (1-45 Hz 3^rd^ order Butterworth bandpass filter; 50 Hz notch 3^rd^ order Butterworth filter; butter and filtfilt functions from Matlab).

### In silico experiments

#### *In silico* model of thalamocortical responses to cortical stimulation

For the *in silico* experiments, we used the model based on our previous works (for model details see^46^. The model included a cortical network with 500 pyramidal neurons (PY) and 100 inhibitory neurons (IN), while the thalamus included 100 thalamocortical neurons (TC) and 100 reticular thalamic neurons (TRN). PY neurons were simulated using a two-compartment model, constituted by a dendritic and a somato-axonal compartment. Conversely, IN, TC, and TRN neurons were simulated using a single-compartment model. The synaptic connectivity between different cell types is given in Fig 4A. Briefly, the connectivity within and between neuron types was implemented by simulating AMPA, NMDA, GABA-A, and GABA-B synapses with local connectivity using a grid-like structure.

For the purposes of this study, we added the simulation of the two conditions: rest and movement The potassium leak currents, synaptic connection strengths and rate of random mini excitatory post synaptic potentials (miniEPSPs) of cortical and thalamic connections were identified for rest condition such that the simulated baseline firing rate was comparable in terms of order of magnitude to baseline firing rate of the mouse *in vivo* resting state data and based on the acetylcholine tone during rest in previous experiments^49–51^. The movement condition was then simulated by increasing miniEPSPs to thalamocortical and reticular neurons by 66.7% (both from 30 mS/cm^2^ to 50 mS/cm^2^) and by reducing the potassium leak current by 26% (acetylcholine level from 1.3 to 1).

We simulated a total of 30 trials of 15 seconds, constituted by 15 rest trials and 15 movement trials. For each trial, the stimulation was administered as a single impulse (duration: 1 ms; intensity: 100 μA) to 50% of the cortical neurons located in the center of the series (i.e. PY neurons 125 to 375; IN neurons 25 to 75). Each pulse was delivered between 8 and 9.75 seconds (with a time difference across trials of 125 ms) from the beginning of the simulation considered to be the time zero.

#### Preprocessing of *in silico* thalamocortical responses

Trials were split in epochs from -2 to +2 seconds around the stimulation onset. Like the mouse *in vivo* analyses, responsive neurons were identified as the neurons with a significant modulation of the intracellular voltage (±5 standard deviations) in the time window between 0 ms and 250 ms post-stimulus onset with respect to the baseline. We then defined two response time windows: an early response window (0-50 ms) and a late response window (75-250 ms). The early time window was used to identify the percentage of responsive trial for each thalamic unit defined as the number of trials with the presence of a spike divided by the total number of trials (i.e. 30 trials, 15 during rest and 15 during movement). The same early time window was used to evaluate the early response latency for cortical neurons (at 0 ms by definition) and for thalamic neurons, calculated as the median of the first spike latency across units. The same computation was applied to the late response window (75-250 ms) to calculate the rebound latencies of cortical and thalamic neurons.

For the late response window, we also evaluated: the evoked thalamic inter-spike interval (ISI), calculated for each TC neuron as the time difference between the first two spikes in the rebound response; the evoked thalamic firing rate, calculated as the inverse of the ISI; and the response onset variability (ROV), assessed for each trial as the standard deviation of the first spike latency across units and then averaged across all trials. Thalamic units were separated into high-firing (HF) and low-firing (LF) units according to the same evoked ISI threshold found in the mice data (i.e. 17.88 ms, Figure 3A).

### Data analysis

#### Evoked potentials (EPs) analyses

Across each mouse experiment, trials were classified by the behavioral state of the animal: quiet wakefulness, if the mouse’s speed (measured by the wheel’s angular velocity) was less than 0.1 cm/s from -0.5 to +0.5 from the stimulus onset; movement, if the mouse’s speed was greater than 0.1 cm/s. For human experiments, rest and movement conditions were a-priori identified by the task performed during each session.

The following analyses were applied on human EEG (ROI contacts), mouse EEG (all contacts), human monopolar and bipolar iEEG (ROI contacts), and mouse LFP and CSD (MOs contacts). The rectified amplitude of the response for each condition was computed as the absolute value of the EPs in the window of interest (early response: 3-50 ms; late response: 150-250 ms). For human and mouse EEG, mouse LFP and mouse CSD recordings, the rectified amplitude for each subject was computed as the average rectified EPs across channels.

The phase locking factor (PLF) for each condition was computed as described by Sinkkonen et al^37^. In brief, the time-resolved phase of each recording site was obtained from the Hilbert transform of each single-trial signal divided by its absolute value and averaged across trials. Then, single-channel PLF was computed by averaging the real part of the phase-vector in the window of interest (early response: 3-50 ms; late response: 150-250 ms). For human and mouse EEG, mouse LFP and mouse CSD recordings, the PLF for each subject was computed as the average PLF across channels.

#### EPs statistical analysis

Movement EPs were compared to rest EPs in terms of both rectified amplitude and PLF of the evoked response. For human EEG, mouse EEG, mouse LFP, and mouse CSD, the average rectified amplitude of the response evoked in each session during movement was compared to the rectified amplitude of the average evoked response at rest through Wilcoxon signed rank test (wilcoxon and stats.wilcoxon functions from Matlab and Scipy – Python, respectively). For human iEEG signals, the squeezing hand task was compared to rest in terms of rectified amplitude and PLF through generalized mixed effects models (lmer function from lmerTest – R; formula: Amplitude ∼ Condition + (1 | SubjectID); PLF ∼ Condition + (1 | SubjectID)). To compare the LFP peak latency in optogenetic experiments, we used a one-way ANOVA (stat.anova_stat from Scipy - Python) and post-hoc comparisons were performed using a Tukey test (stat. tukey_hsd from Scipy - Python).

#### Unit analyses

The instantaneous firing rate response of each unit was computed through an adaptive kernel algorithm for firing estimation^123^. Units were considered responsive if their response to the stimulation (3 to 500 ms from stimulation onset) exceeded ±7 standard deviations of the baseline (- 500 to -50 ms) firing rate. Average firing rates were computed as the average number of spikes in the time window of interest for each trial divided by the time window length.

The latency of units’ response was quantified as the median latency of the first spike in the window of interest across trials. The evoked response onset variability (ROV) was assessed as the standard deviation of the first spike latency across units for each trial.

The evoked inter-spike-interval (ISI) was quantified as the median across trials of the latency between the first spike and the subsequent spike in the time window of interest. The evoked response frequency was assessed as the reciprocal of the evoked inter-spike-interval (ISI) in the time window of interest (i.e. early response: 3-50 ms; late response: 150-250 ms).

Cross-correlograms were computed by binning (bin size = 2 ms) the absolute value of the triangular matrix of the delay between each spike for each pair of neurons divided by the number of seed spikes. Statistical tests were computed on the cross-correlation values obtained as the average of the instantaneous cross-correlation over the first 10 ms (i.e. 5 bins).

#### Unit statistical analyses

The movement condition was compared to rest in terms of units evoked firing rate through a Wilcoxon signed rank test (stats.wilcoxon function from Scipy - Python). Putative thalamic relay units were separated into high-firing (HF) and low-firing (LF) units through a data-driven cluster analysis (cluster.KMeans from sklearn - Python) of the ISI of the evoked response in the late time window (150-250 ms from stimulation onset). Comparison between rest and movement condition for HF and LF units in terms of baseline firing rate, response frequency, and response onset variability was performed through Wilcoxon signed rank tests (stats.wilcoxon function from Scipy - Python) and Bonferroni corrected for multiple comparison (stats.multitest.multipletests function from statsmodels - Python). The statistical comparison between cross-correlation values was computed through Wilcoxon signed rank tests (stats.wilcoxon function from Scipy - Python).

Correlation of baseline firing rate with response frequency and response onset variability for HF and LF units was performed through generalized mixed effects models (smf.mixedlm function from statmodels - Python; Formula: UnitFeature ∼ BaselineFr + (1 | MouseID)). Similarly, correlation of response frequency and response onset variability for HF and LF units with evoked EEG rectified amplitude was performed through a mixed effect model (smf.mixedlm function from statmodels - Python; Formula: EEG Amplitude ∼ ResponseFrequencyHF + ResponseVariabilityHF + ResponseFrequencyLF + ResponseVariabilityLF + (1 | MouseID)).

