## Supplemental materials for "Thalamic feedback shapes brain responses evoked by cortical stimulation in mice and humans"

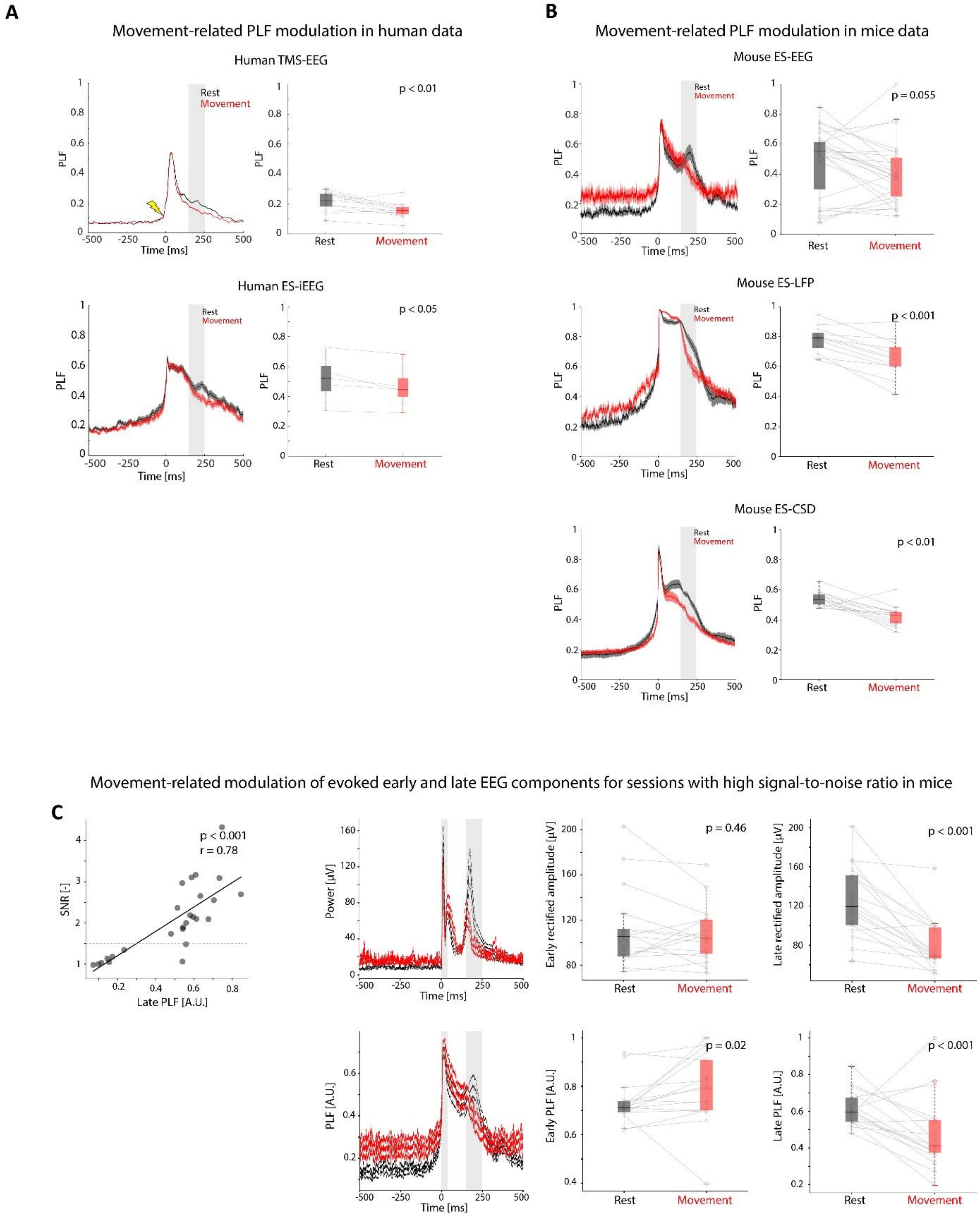

**Figure S1. Behavioral-state-dependent modulation of the phase-locking factor (PLF) in humans and mice.**

A) PLF as a function of time from the stimulation onset (left) and averaged PLF calculated over the late response time window (150-250 ms, shaded grey) at rest (black) and movement (red) for human TMS-EEG (top) and human ES-iEEG (bottom). The average PLF of the late response evoked by TMS in humans decreased from 0.21±0.07 during rest to 0.15±0.05 during movement (Wilcoxon signed rank test, p=0.002). The average PLF of the response evoked by ES in humans decreased from 0.55±0.27 during rest to 0.49±0.27 during movement (mixed-effect model ES-iEEG p=0.033).

B) PLF as a function of time from the stimulation onset and averaged PLF calculated over the late response time window (150-250 ms, shaded grey) at rest (black) and movement (red) in mice across recording modalities: EEG, LFP, CSD (Wilcoxon signed rank test: ES-EEG p=0.055; ES-LFP p<0.001; ES-CSD p<0.01).

C) EEG modulation of early and late components (3-50 ms and 150-250 ms, respectively; shaded grey) in sessions with high signal-to-noise ratio (SNR) in mice at rest and during movement. Left: Relationship between SNR and PLF calculated for the late component (Spearman correlation, p<0.001, r = 0.78). Right panels: modulation of the EEG evoked rectified amplitude (top) and PLF (bottom) of early and late component only for the EEG sessions with high SNR (SNR > 1.5; Amplitude early: p=0.46; Amplitude late: p<0.001; PLF early: p=0.02; PLF late: p<0.001).

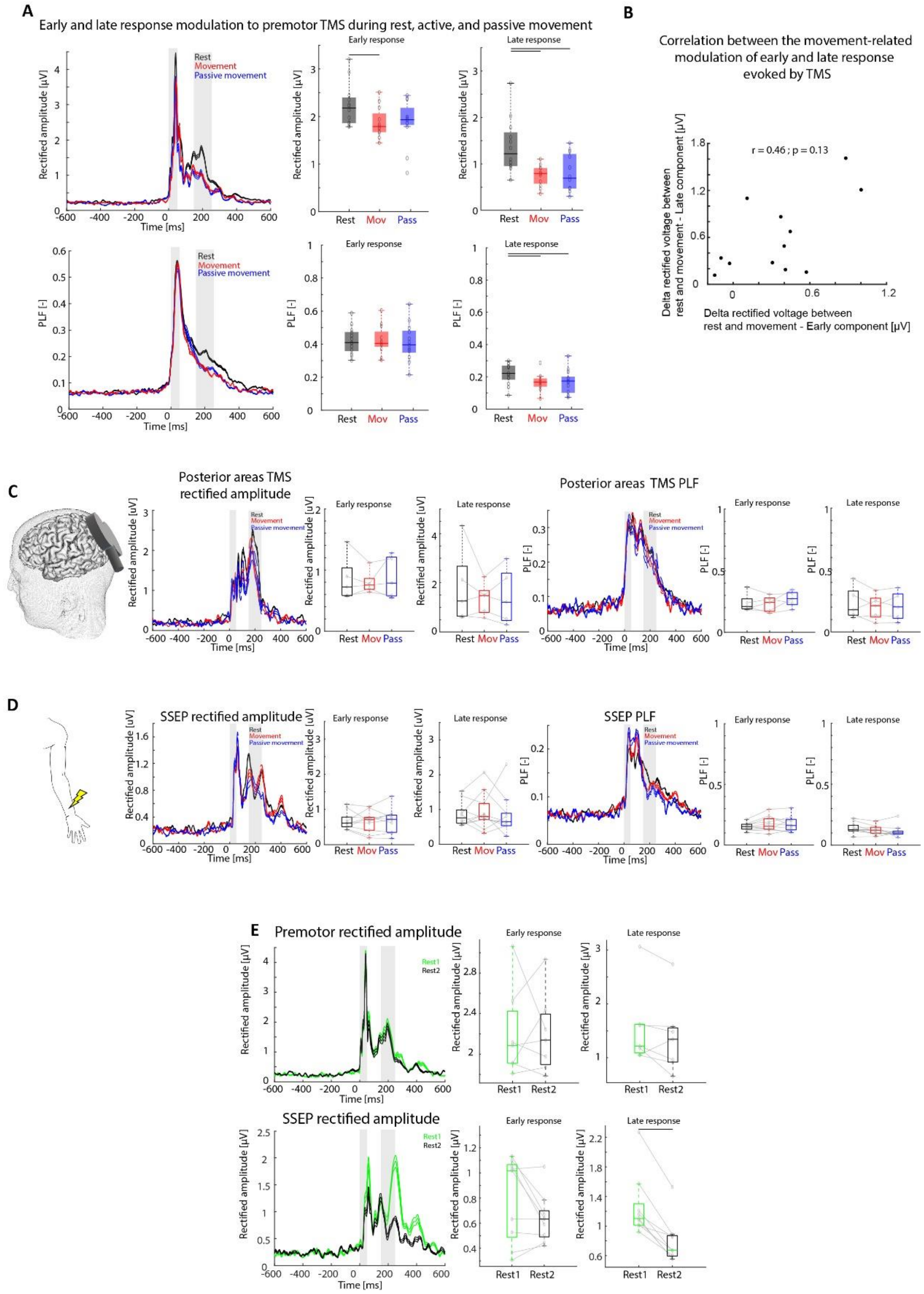


**Figure S2. Controls for the TMS-EEG experiments in humans.**

A) Modulation of the EEG responses to TMS delivered to premotor cortex in humans at rest (black), during active (red) and passive (blue) movement. Top row: Premotor evoked EEG rectified amplitude over time during rest and movement. EEG rectified amplitude average calculated over the early response time window (3-50 ms, shaded grey left), and over the late response time window (150-250 ms, shaded grey left, right) at rest and during active and passive movement for each subject (*Wilcoxon signed rank test; rest VS passive: Earl:*

*p=0.068, ns; Late: p<0.01; rest VS active: Early: p<0.01; Late: p<0.01; active VS passive: Early: p = 0.56; Late: p=1*). Bottom row: Premotor evoked EEG phase locking factor (PLF) over time during rest and movement. EEG PLF calculated over the early (3-50 ms, shaded grey, left) and late (150-250 ms, shaded grey, right) response time windows during rest and passive movement for each subject (voltage at rest 2.24±0.45 μV versus 1.88±0.32 μV during movement; PLF at rest: 0.43±0.8 versus movement: 0.43±0.9; *Wilcoxon signed rank test; rest VS passive: Early p=0.94, ns; Late p<0.01; rest VS active: Early: 0.97; Late: p<0.01; active VS passive: Early: p=0.47; Late: p=0.79*).

B) Lack of correlation between the TMS early and late components modulated by movement (n=12 subjects; Pearson's correlation; r=0.46; p=0.13, ns).

C) Lack of modulation of EEG responses to TMS delivered to occipital cortical areas in humans at rest (black), during active (red) and passive (blue) movement. From left to right. Schematics of the setup with a TMS coil stimulating the occipital area. Evoked EEG rectified amplitude over time during rest, active and passive movement. EEG rectified amplitude average calculated over the early response time window (3-50 ms, shaded grey) for the three conditions (*Wilcoxon signed rank tests; rest-movement: p=0.56, ns; rest-passive: p=1, ns*). EEG rectified amplitude average calculated over the late response time window (150-250 ms, shaded grey) for the three conditions (*Wilcoxon signed rank tests; rest-movement: p=0.31, ns; rest-passive: p=0.31, ns*). Evoked EEG phase locking factor (PLF) over time for the three conditions. EEG PLF calculated over the early (3-50 ms, shaded grey) and late (150-250 ms, shaded grey) response time windows for each subject (*Wilcoxon signed rank tests; early and late responses rest-movement: p=0.84, ns; p=0.56, ns; early and late responses rest-passive: p=0.81, ns; p=0.44, ns*).

D) Lack of modulation of EEG responses to electrical stimulation of the left median nerve (90% motor threshold) at rest (black), active (red) and passive (blue) movement. The plots are structured in the same fashion as C (*Wilcoxon signed rank tests; EEG rectified amplitude early rest-movement: p=0.41, ns; rest-passive: p=0.92, ns; EEG rectified amplitude late rest-movement: p=0.96, ns; rest-passive: p=0.37, ns; EEG PLF early rest-movement: p= 0.21, ns; rest-passive: p= 0.16, ns; EEG PLF late rest-movement: p= 0.17, ns; rest-passive: p= 0.19, ns*).

E) Adaptation over time of the late EEG response to the somatosensory stimulation of the median nerve, as opposed to a lack of adaptation for the TMS evoked responses in humans. Top: comparison of EEG responses to TMS of premotor cortex in humans between early and late sessions at rest. From left to right. Evoked EEG rectified amplitude over time during the first (green) and second (black) sessions at rest. EEG rectified amplitude average calculated over the early response time window (3-50 ms, shaded grey, left) and over the late response time window (150-250 ms, shaded grey, right) for the two conditions *(Wilcoxon signed rank tests; Early p= 1, ns; Late p= 0.46, ns*). Bottom: Comparison of somatosensory evoked responses (SSEP) evoked by electrical stimulation of the left median nerve between early and late sessions at rest. Plots are structured in the same fashion as above (*Wilcoxon signed rank tests; Early p= 0.43, ns; Late p<0.01*).

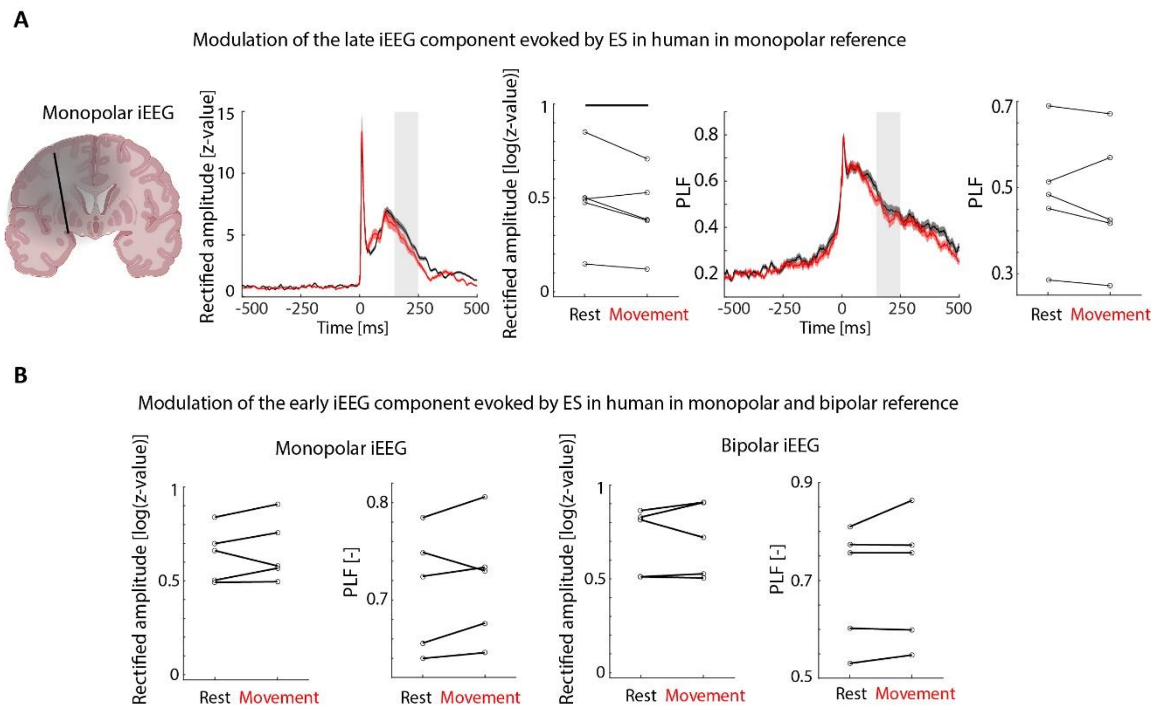


**Figure S3. Monopolar derivation of the iEEG recording and associated quantifications.**

A) Modulation of monopolar iEEG evoked responses in humans at rest and during movement. Similar visualization shown in Figure 1B, but for monopolar iEEG configuration (154 contacts from 5 subjects*;* Rectified amplitude rest: 3.59±3.45 z-value; Rectified amplitude movement:5.8±5.32 z-value; PLF Rest: 0.42±0.22; PLF Movement: 0.47±0.26; *Linear Mixed Effect Models; Rectified amplitude: p=0.037; PLF: p=0.27, ns*).

B) Modulation of the early iEEG response evoked by electrically stimulating supplementary motor cortex in human at rest and during movement. Left: monopolar iEEG rectified amplitude (left) and PLF (right) average over the early response window (3-50 ms) at rest and movement for all ROI contacts across subjects (*Mixed effect model; lmer function from lmerTest - R; Rectified amplitude: p = 0.91, ns; PLF: p = 0.93, ns*). Right: bipolar iEEG rectified amplitude (left) and PLF (right) average over the early response window (3-50 ms) at rest and movement for all ROI contacts across subjects (*Mixed effect model; lmer function from lmerTest - R; Rectified amplitude: p = 0.99, ns; PLF: p = 0.88, ns*).

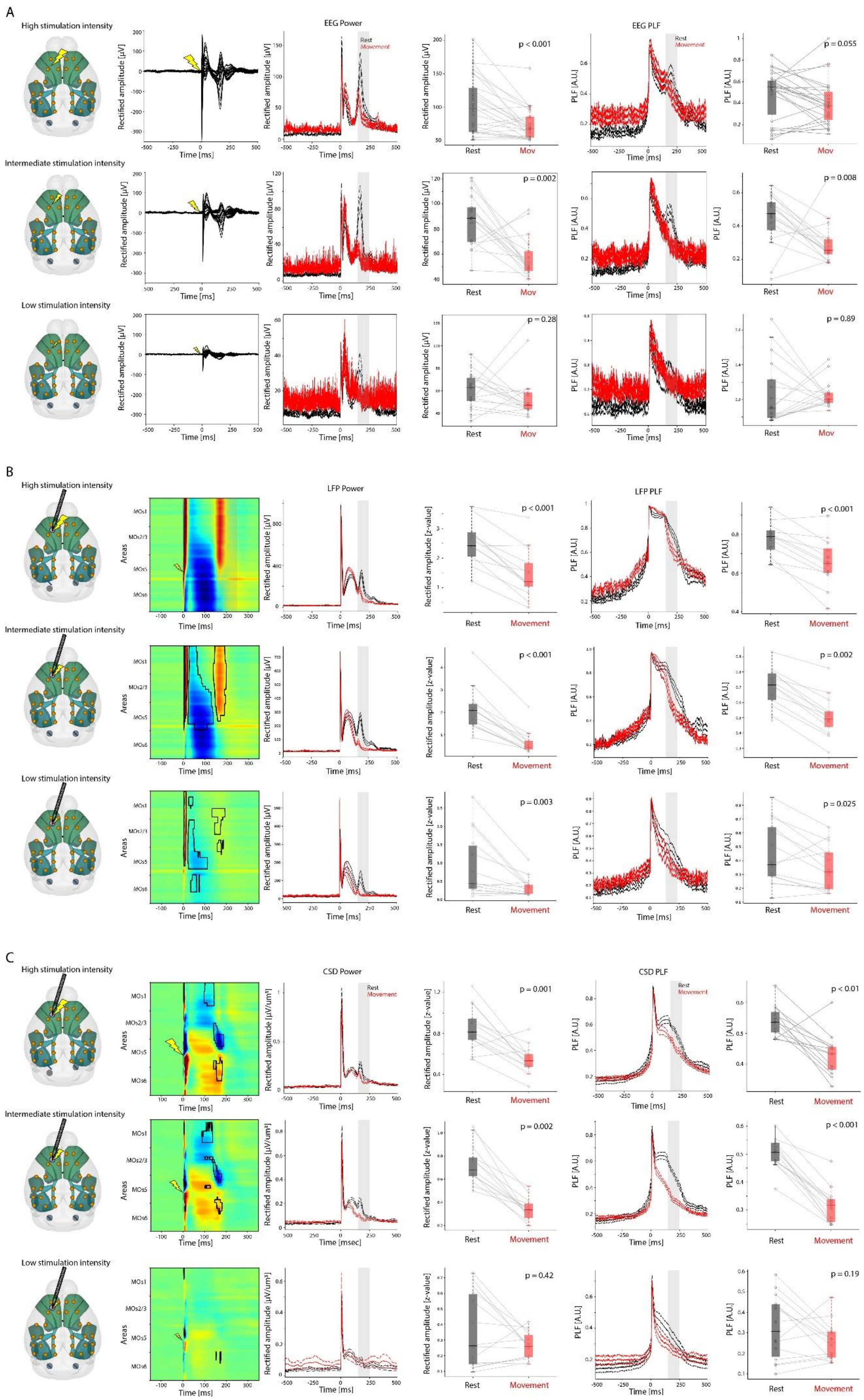
A
High stimulation intensity
Intermediate stimulation intensity
Low stimulation intensity
EEG Power
EEG PLF
Rest
Movement
Rectified amplitude [µV]
PLF [A.U.]
Time [ms]
p < 0.001
p = 0.055
p = 0.002
p = 0.008
p = 0.28
p = 0.89
Mov
B
LFP Power
LFP PLF
Areas
MOs1
MOs2/3
MOs5
MOs6
Rectified amplitude [z-value]
p < 0.001
p = 0.002
p = 0.003
p = 0.025
C
CSD Power
CSD PLF
Rectified amplitude [µV/um³]
p = 0.001
p < 0.01
p < 0.001
p = 0.42
p = 0.19
Time [msec]

**Figure S4. Modulation of the late response evoked by electrical stimulation of MOs in mice as a function of current intensities at rest and during movement.**

A) Modulation of the EEG response to the electrical stimulation of MOs in mice during rest and movement across stimulation current intensities. From top to bottom, highest to lowest stimulation intensity (High: 66.15±12.11 uA; Intermediate: 47.35±13.30 uA; Low: 30.59±15.80 uA). From left to right. Schematics of the EEG recording array and the location relative to it of the stimulated area; Butterfly plot of ES-evoked responses (-500 to 500 ms around stimulus onset) for all EEG electrode traces across all subjects (n=21). Grand-average (21 subjects) during rest (black) and movement (red). The rectified amplitude average of the late component (150-250 ms; shaded grey) during rest and movement for each subject (*Wilcoxon signed rank test; High: $p < 0.001$; Intermediate: $p = 0.002$; Low: $p = 0.28$, ns*). EEG phase locking factor (PLF) evoked at rest (black) and movement (red) calculated over all EEG contacts and subjects. EEG PLF averaged over the late response window (150-250 ms, shaded grey) at rest and movement across subjects (*Wilcoxon signed rank test; High: $p = 0.055$, ns; Intermediate: $p = 0.008$; Low: $p = 0.89$, ns*).

B) Modulation of the local field potential (LFP) responses from the Neuropixels probes in MOs evoked by the ES of the deep layers of the same area at rest and during movement. Plots are structured in the same fashion as in A, with the voltage displayed in a color map as a function of depth from superficial to deep layers represented in y-axis from top to bottom (*Wilcoxon signed rank test; Rectified amplitude high: $p < 0.001$; Rectified amplitude intermediate: $p < 0.001$; Rectified amplitude low: $p = 0.003$; PLF high: $p < 0.001$; PLF intermediate: $p =0.002$; PLF low: $p = 0.025$*).

C) Modulation of the current source density (CSD) estimated from the Neuropixels in MOs where the electrical stimulation was delivered at rest and during movement. Same structure as B (*Wilcoxon signed rank test; Rectified amplitude high: $p < 0.001$; Rectified amplitude intermediate: $p = 0.002$; Rectified amplitude low: $p = 0.42$, ns; PLF high: $p < 0.01$; PLF intermediate: $p < 0.001$; PLF low: $p = 0.19$, ns*). Notice that grand-average CSD at high and intermediate stimulation intensities shows a sink at the border between layer 2/3 and 5 - suggesting an input from the thalamus. The same sink is no longer visible when low current intensity is applied.

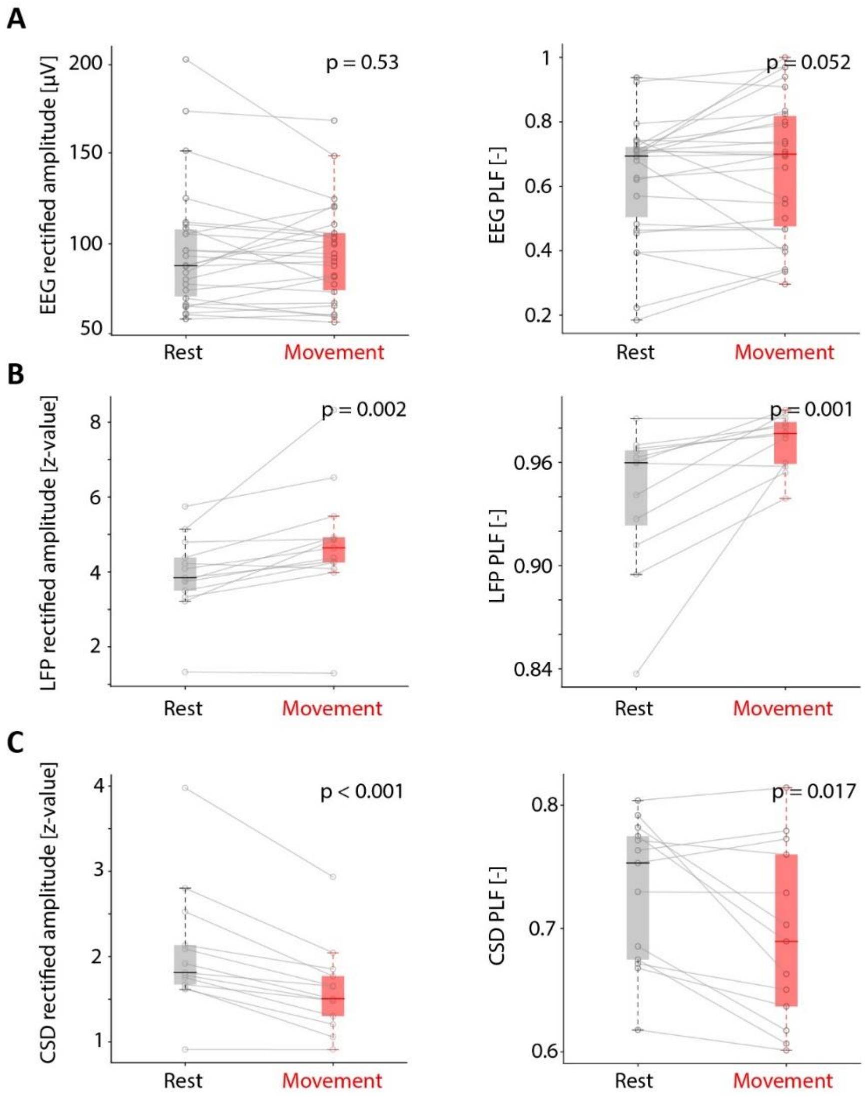


**Figure S5. Modulation of the early component evoked by the electrical stimulation of MOs in mice at rest and during movement.**

A) Absence of modulation for the early EEG response to MOs electrical stimulation in mice (n=15) at rest and during movement. Left panel: EEG rectified amplitude averaged over the early response window (3-50 ms) at rest and movement across subjects. (*Wilcoxon signed rank test; p = 0.53, ns*). EEG PLF averaged over the early response window (3-50 ms) at rest and movement across subjects (*Wilcoxon signed rank test; p = 0.052, ns*).

B) Early LFP responses from the Neuropixels probes in MOs evoked by the ES of the deep layers of the same area at rest and movement across subjects (n=13). Plots are structured in the same fashion as in A (*Wilcoxon signed rank test; Rectified amplitude: p=0.002; PLF: p = 0.001, movement larger than rest*).

C) Early response of the current source density (CSD) estimated from the Neuropixels in MOs at rest and movement across subjects. Plots are structured in the same fashion as in A and B (*Wilcoxon signed rank test; Rectified amplitude: p<0.001; PLF: p = 0.017*).

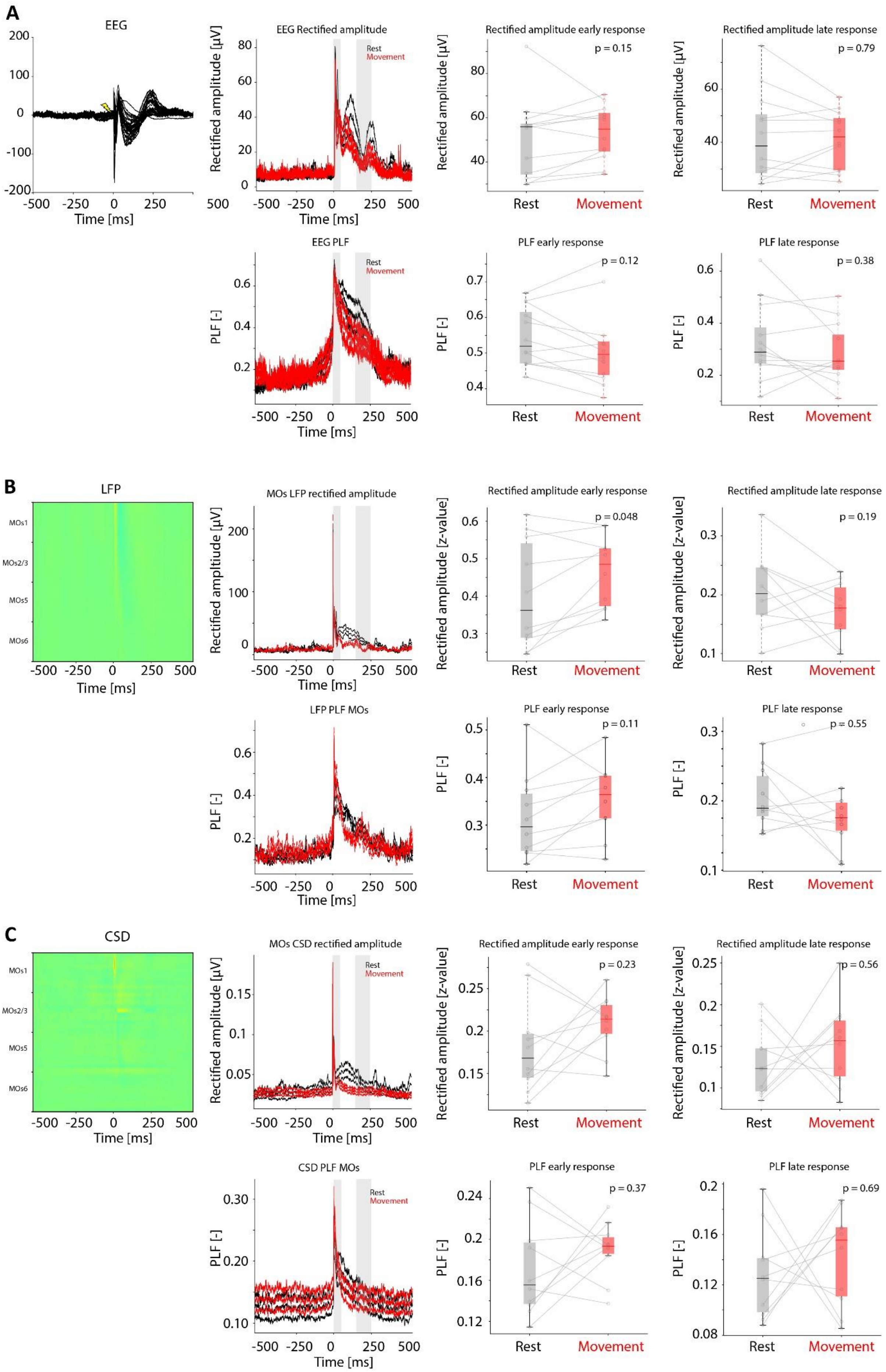
A
EEG
Time [ms]
EEG Rectified amplitude
Rest
Movement
Rectified amplitude [µV]
Rectified amplitude early response
p = 0.15
Rectified amplitude late response
p = 0.79
EEG PLF
PLF [-]
PLF early response
p = 0.12
PLF late response
p = 0.38
Rest
Movement
B
LFP
MOs1
MOs2/3
MOs5
MOs6
MOs LFP rectified amplitude
Rectified amplitude early response
p = 0.048
Rectified amplitude [z-value]
Rectified amplitude late response
p = 0.19
LFP PLF MOs
PLF early response
p = 0.11
PLF late response
p = 0.55
C
CSD
MOs CSD rectified amplitude
Rectified amplitude early response
p = 0.23
Rectified amplitude late response
p = 0.56
CSD PLF MOs
PLF early response
p = 0.37
PLF late response
p = 0.69

**Figure S6. Lack of modulation of both early and late responses across recording modalities evoked by electrical stimulation of posterior cortical areas in mice at rest and during movement.**

A) Absence of modulation of the EEG response to the electrical stimulation of control areas (visual cortex, retrosplenial cortex, hippocampus) in mice during rest (black) and movement (red). From left to right: Evoked potentials (EPs; -500 to 500 ms around stimulus onset) across subjects (n = 12 sessions from 6 subjects) for all EEG electrode traces superimposed (butterfly plot); EEG rectified amplitude evoked at rest (black) and movement (red) calculated over all EEG contacts for all subjects; EEG rectified amplitude averaged over the early response window (3-50 ms) at rest and movement across subjects (*Wilcoxon signed rank test; p = 0.15, ns*). EEG rectified amplitude averaged over the late response window (150-250 ms) at rest and movement across subjects (*Wilcoxon signed rank test; p = 0.79, ns*). EEG phase locking factor (PLF) evoked at rest (black) and movement (red) calculated over all EEG contacts and subjects. EEG PLF averaged over the early response window (3-50 ms) at rest and movement across subjects (*Wilcoxon signed rank test; p = 0.12, ns*). EEG PLF averaged over the late response window (150-250 ms) at rest and movement across subjects (*Wilcoxon signed rank test; p = 0.38, ns*).

B) Absence of modulation in the LFP responses of the Neuropixels in MOs evoked by the electrical stimulation of control areas in mice during rest and movement (n = 10 sessions from 5 subjects). Plots are structured in the same fashion as in A, with the voltage displayed in a color map as a function of depth from surficial to deep layers represented on y-axis from top to bottom (*Wilcoxon signed rank tests; Early rectified amplitude: p = 0.048; Early PLF: p = 0.19, ns; Late rectified amplitude: p = 0.11, ns; Late PLF: p=0.55*).

C) Absence of modulation of the CSD estimated from the LFPs of the Neuropixels in MOs evoked by the electrical stimulation of control areas in mice during rest and movement (n = 10 sessions from 5 subjects). Plots are structured in the same fashion as in B (*Wilcoxon signed rank tests; Early rectified amplitude: p = 0.23, ns; Early PLF: p = 0.56, ns; Late rectified amplitude: p = 0.37, ns; Late PLF: p = 0.69, ns*). Notice that, as opposed to the CSD response evoked by MOs stimulation (Figure 1, S4), these CSD plots do not show any sink in between layers 2/3 and 5.

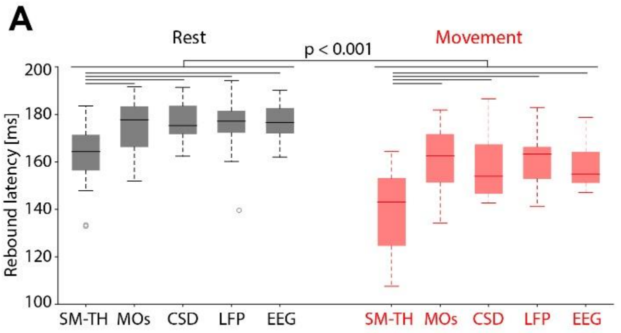


| | Rest | Movement |
|---|---|---|
| EEG | 176 ± 7.2 ms | 158 ± 9.1 ms |
| LFP | 175 ± 11.2 ms | 161 ± 9.9 ms |
| CSD | 177 ± 8.6 ms | 158 ± 13.3 ms |
| MOs units | 176 ± 10.1 ms | 162 ± 12.3 ms |
| SM-TH units | 163 ± 13.1 ms | 139 ± 16.4 ms |

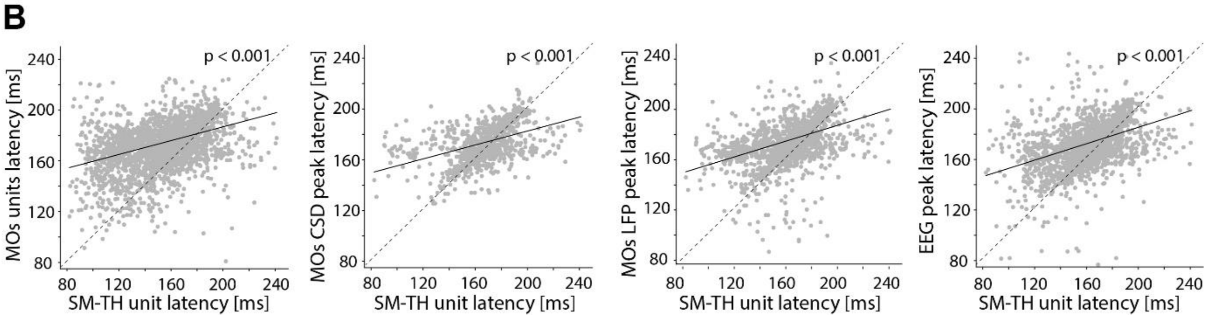


**Figure S7. Latencies of the rebound responses at rest and movement across recording modalities to electrical stimulation of MOs in mice.**

A) Timing of the late responses for rest (black) and movement (red) across different electrophysiological signals to MOs electrical stimulation. Global mean latencies of SM-TH and MOs units (calculated for each subject as the median across units of the first spike in the late response window and then averaged across subjects) and latencies of the CSD, LFP, and EEG peaks at rest (black) and movement (red). Responses during movement showed shorter latencies than at rest for all the evaluated matrices (as previously reported (Claar et al 2023), with SM-TH units preceding any other response (two-way ANOVA, Tukey post-hoc test; $p<0.001$ for all significance lines shown in the figure; $p>0.05$, ns for all the other comparisons). The table to the right reports the exact latency values in milliseconds for each recording modality (mean ± standard deviation).

B) Late response's latencies of MOs units (1395 cortical units), CSD peak, LFP peak, and EEG peak as a function of the latency of the first thalamic spike (1269 thalamic units) in the same time window across all subjects (n=21). All these measurements show a significant positive correlation. Specifically, we found that the latency of thalamic units was positively correlated with the latency of cortical units (First panel, *Mixed effect model;* Formula: *CorticalUnits ~ ThalamicUnits + (1 | MouseID); p <0.001*), as well as with the latency of the CSD peak (Second panel, *Mixed effect model;* Formula: *CSDPeak ~ ThalamicUnits + (1 | MouseID); p <0.001*), with the latency of the LFP peak (Third panel, *Mixed effect model;* Formula: *LFPPeak~ ThalamicUnits + (1 | MouseID); p <0.001*) and with the latency of the EEG peak (Fourth panel, *Mixed effect model;* Formula: *EEGPeak ~ ThalamicUnits + (1 | MouseID); p <0.001*). Each point in this scatter-diagram corresponds to one trial in one mouse.

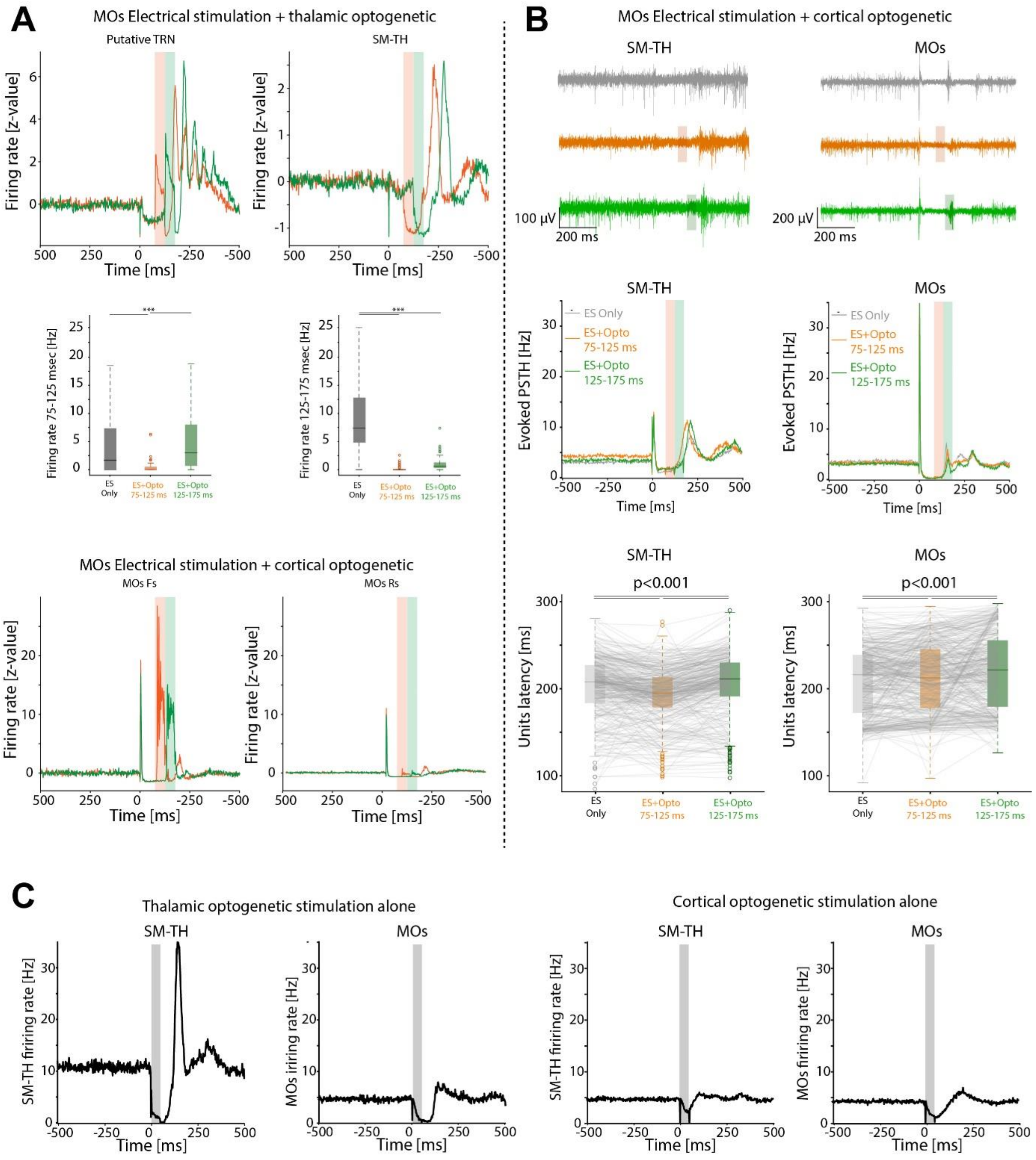


**D**

| | Rest | | Movement | |
|---|---|---|---|---|
| | SM-TH | MOs | SM-TH | MOs |
| ES Only | 166±33 | 199±32 | 151±37 | 202±40 |
| ES + Opto 75-125 | 215±22 | 209±27 | 211±24 | 214±30 |
| ES + Opto 125-175 | 216±60 | 227±28 | 204±68 | 250±29 |

**Figure S8. Characterization of the neural responses to optogenetic stimulation of GABAergic neurons in thalamus and cortex in VGAT-ChR2-YFP/wt mice.**

A) Top: normalized peristimulus time histogram of TRN units (left) and relay thalamic units (right) in one representative mouse during early (75-125 ms, orange) and late optogenetic activation (125-175 ms, green). TRN units exhibit an abrupt increase in the firing rate during optogenetic activation while the relay thalamic units abruptly decrease firing rate in the same time windows. Middle: average firing rate of putative relay thalamic units in the 75-125 ms window (left) and in the 125-175 ms window (right) during electrical stimulation alone (black), electrical stimulation and thalamic opto-inhibition in the 75-125 ms window (orange), and during electrical stimulation and thalamic opto-inhibition in the 125-175 ms window (green). The activity in the 75-125 window is significantly reduced by thalamic opto inhibition in the same window ($p<0.001$; one-way ANOVA). Conversely, the activity in the 125-175 window is significantly reduced by both optogenetic protocols ($p<0.001$; one-way ANOVA). Bottom: normalized peristimulus time histogram of putative inhibitory cortical units (FS, left) and putative excitatory units (RS, right) in one representative mouse during early (75-125 ms, orange) and late optogenetic activation (125-175 ms, green) of cortical GABAergic neurons.

B) Cortical optogenetic inhibition paired with electrical stimulation induced a minimal delay compared to thalamic inhibition (Figure 2) of the cortical rebound response to electrical stimulation. Top: representative traces of the action potential band in SM-TH and MOs RS evoked by electrical stimulation (grey), MOs optogenetic stimulation delivered between 75 and 125 ms after electrical stimulation (orange), and MOs optogenetic stimulation delivered between 125 and 175 ms after electrical stimulation (green). Middle: peristimulus time histogram (PSTH) for 361 SM-TH units' and 417 MOs units' (RS) responses elicited by the electrical stimulation combined with MOs optogenetic stimulation averaged across 2 mice. Color code as above. Bottom: units' latencies of the cortical evoked responses plotted above and elicited by the same stimulation protocols (Wilcoxon signed rank tests; all $p<0.001$; MOs: ES Only=208±39 ms; ES+Opto75-125=213±40 ms; ES+Opto125-175=220±44 ms; SM-TH: ES Only=201±36 ms; ES+Opto75-125=191±32 ms; ES+Opto125-175=204±36 ms).

C) Optogenetic stimulation alone of GABAergic neurons in cortex and thalamus. Left: average firing rate of SM-TH units (129 units, left) and MOs units (156 units, right) evoked by optogenetic activation of thalamic GABAergic neurons strongly decreases firing rate of SM-TH and MOs units (grey shaded area highlights the optogenetic stimulation). The thalamic suppression is followed by a large rebound excitation. Right: average firing rate of SM-TH units (417 units, left) and MOs units (440 units, right) evoked by optogenetic activation of MOs GABAergic neurons showing a strong decrease in firing rate for both SM-TH and MOs units.

D) Latencies of SM-TH and MOs rebound responses following electrical stimulation of MOs and optogenetic suppression of the thalamus in mice during rest and movement.

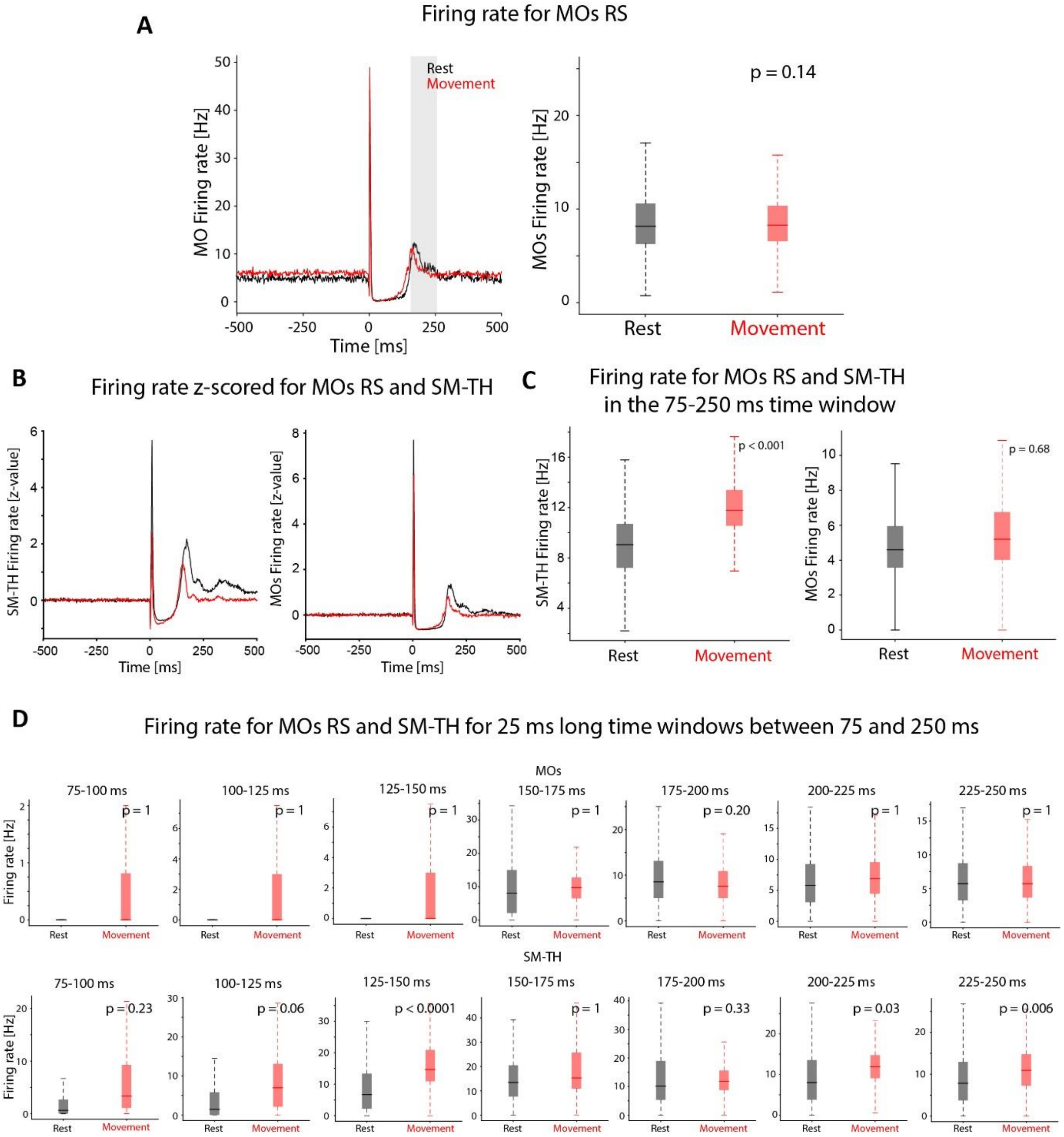


**Figure S9. Lack of modulation of MOs and SM-TH firing rate at rest and during movement.**

A) Lack of modulation of MOs units' evoked responses. Left: Evoked MOs firing rate at the population level during rest and movement (black and red, respectively; 1009 units). Right: MOs firing rate averaged over the late response (150-250 ms, shaded grey) during rest and movement across subjects (n=19 subjects*, Wilcoxon signed rank test p = 0.14, ns*).

B) Modulation of the normalized firing rate, reported as z-score of the average pre-stimulus onset firing, of all the neurons for RS neurons in SM-TH (left) and MOs (right) at rest and during movement.

C) Comparison between rest (black) and movement (red) of the evoked firing rate over an extended time window between 75 ms and 250 ms from stimulation onset for RS neurons in SM-TH (left; Wilcoxon rank sum test, p <0.001) and MOs (right; Wilcoxon rank sum test, p = 0.68, ns).

D) Comparison of MOs (top) and SM-TH (bottom) firing rate evoked by electrical stimulation of MOs between rest (black) and movement (red) calculated over 25 ms long windows from 75 ms to 250 ms after stimulus onset.

C and D demonstrate that throughout the entire response, the evoked firing rate at rest never exceeds evoked firing rate during movement (Wilcoxon rank sum test, Benjamini-Yekutieli false-discovery-rate correction for multiple comparison (Benjamini-Yekutieli, Ann. Statist. 2001); *fdr_BY* function from *Multiple Testing Toolbox* - Matlab).

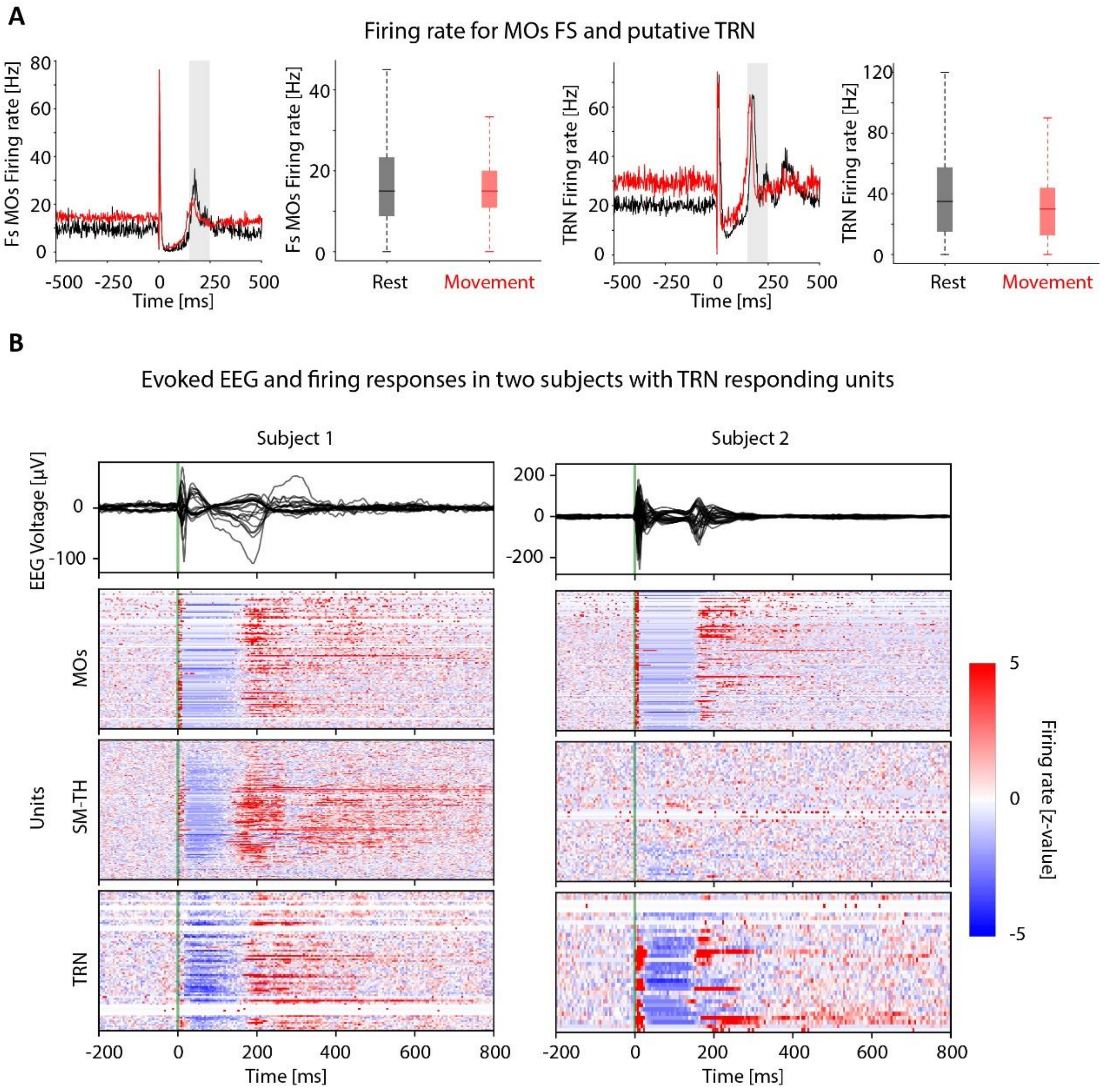


**Figure S10. Electrically evoked responses of cortical and thalamic GABAergic neurons.**

A) Modulation of putative inhibitory cortical neurons (FS) and putative TRN inhibitory neurons at rest and during movement. Evoked firing rate at the population level at rest and movement (black and red, respectively) for MOs FS units (145 units from 19 mice) and putative TRN neurons (67 units from 21 mice) with associated quantification over the late response window (*Wilcoxon signed rank test; FS p = 0.35, ns; TRN p = 0.77, ns*).

B) Recordings from two subjects (left and right columns) showing the responses to MOs electrical stimulation evoked in terms of EEG (top panel) and spiking activity expressed as normalized firing rate, reported as z-score of the average pre-stimulus firing rate for all units in MOs, SM-TH, and TRN (bottom panels).

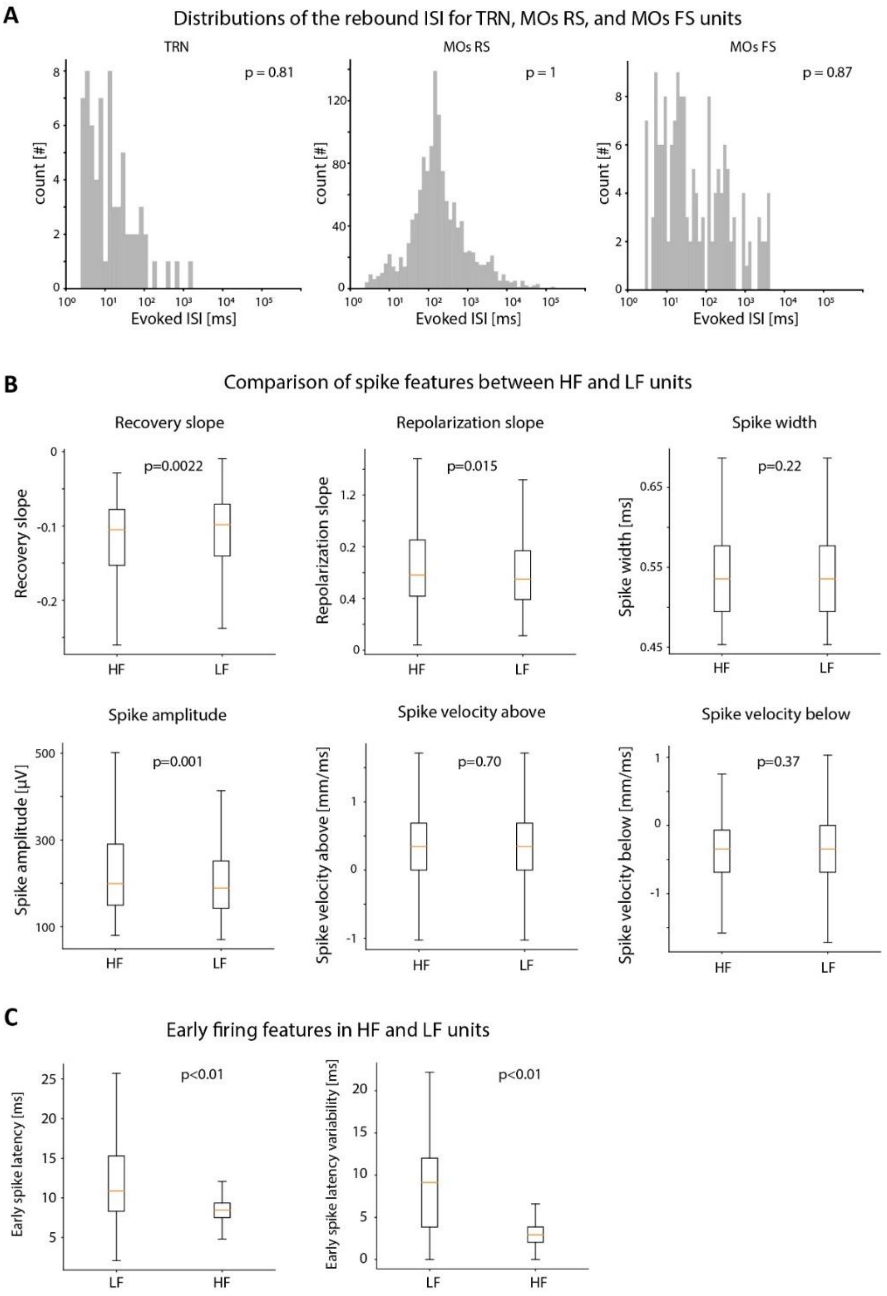


**S11. Characterization of thalamic units' responses.**

A) Evoked ISI histogram in the late response to MOs stimulation for the different putative neuronal populations: thalamic TRN and cortical (RS and FS). Hartigans' dip test indicates that none of these distributions are bimodally distributed (TRN $p = 0.81$; MOs RS $p=1$; MOs FS $p = 0.87$), as opposed to the putative relay thalamic units shown in Figure 3C.

B) Comparison of morphological spike features extracted by Kilosort (Stringer et al., 2019) between HF and LF units in mice. No major differences were found between HF and LF suggesting that they don't belong to different cell types.

C) Early response latency (*Wilcoxon signed rank test; $p < 0.01$*) and latency variability (*Wilcoxon signed rank test; $p < 0.01$*) of thalamic LF and HF units evoked by MOs electrical stimulation.

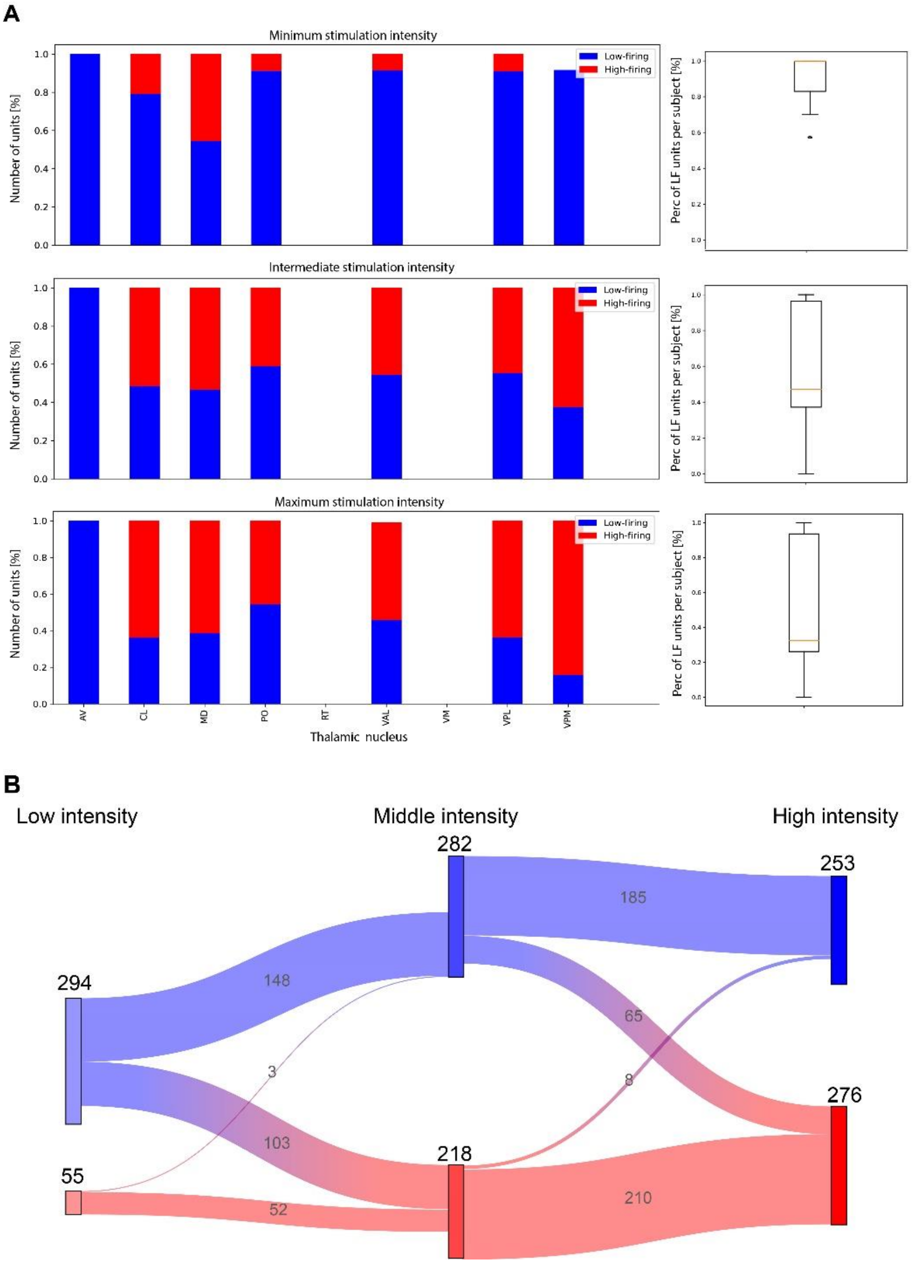


**Figure S12. Current intensity modulates the relative percentage of thalamic units classified as LF and HF.**

A) Percentage of units classified as HF (red) and LF (blue) for low, intermediate, and high current intensities across thalamic nuclei (left). Averaged percentage of thalamic units classified as LF across thalamic nuclei for all the subjects (right).

B) Sankey diagram showing the number of units classified as HF (red) and LF (blue) for low, intermediate, and high current intensities. The diagram demonstrates that LF classified units progressively become HF classified units as current intensity increases.

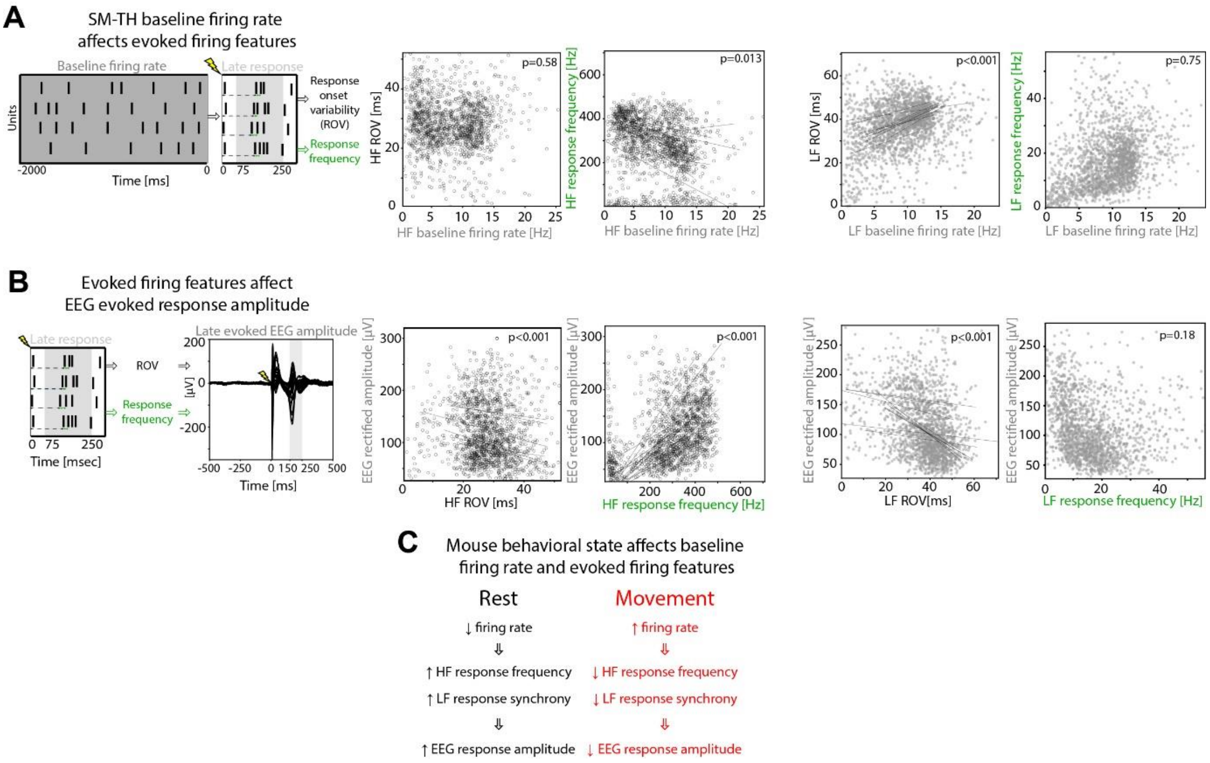


**S13. Modulation of the late EP component evoked by electrical stimulation correlates with the modulation of thalamic unit's synchrony in mice.**

A) Relation between thalamic baseline firing rate, response onset variability (ROV), and evoked response frequency. ROV across units for each trial as a function of average baseline firing rates for HF units across subjects (no significant correlation: *Linear mixed effect model*, p=0.58, ns). Late evoked response frequency as a function of baseline firing rates for HF units across trials and subjects (*Linear mixed effect model, p=0.012*). Plots on the right are structed in the same fashion as the ones on the left, but for LF units (*Linear mixed effect models; Baseline firing rate-Evoked response variability: p<0.001; Baseline firing rate-Evoked response frequency:* p=0.75, ns).

B) Relation between thalamic units' response features and the magnitude of the EEG evoked late responses. From left to right: schematic representation of the considered thalamic units' response features (i.e. response onset variability [ROV] and evoked response frequency) and the extracted EEG magnitude (calculated as the EEG rectified amplitude of the late evoked responses; 150-250 ms, grey shaded). Evoked EEG rectified amplitude of the late response as function of late evoked ROV (*Linear mixed effect model, p<0.001*) and evoked response frequency for HF units (*Linear mixed effect model, p<0.001*). Plots on the right are structed in the same fashion as the ones on the left, but for LF units (*Linear mixed effect model; Evoked response variability-EEG rectified amplitude: p<0.001; Evoked response frequency-EEG rectified amplitude:* p=0.18, ns).

C) Summary of the state-dependent modulation of the electrically evoked EEG responses shown in D and E: compared to movement, during resting, the thalamus has a lower spontaneous firing rate; this reduction in the thalamic firing rate is associated with an increase of the HF units response frequency and an increase of LF units synchronicity (i.e. decrease of ROV) in the late response. As a result of these modulations at the thalamic units' level, the rectified amplitude of the late EEG response is larger at rest compared to movement.

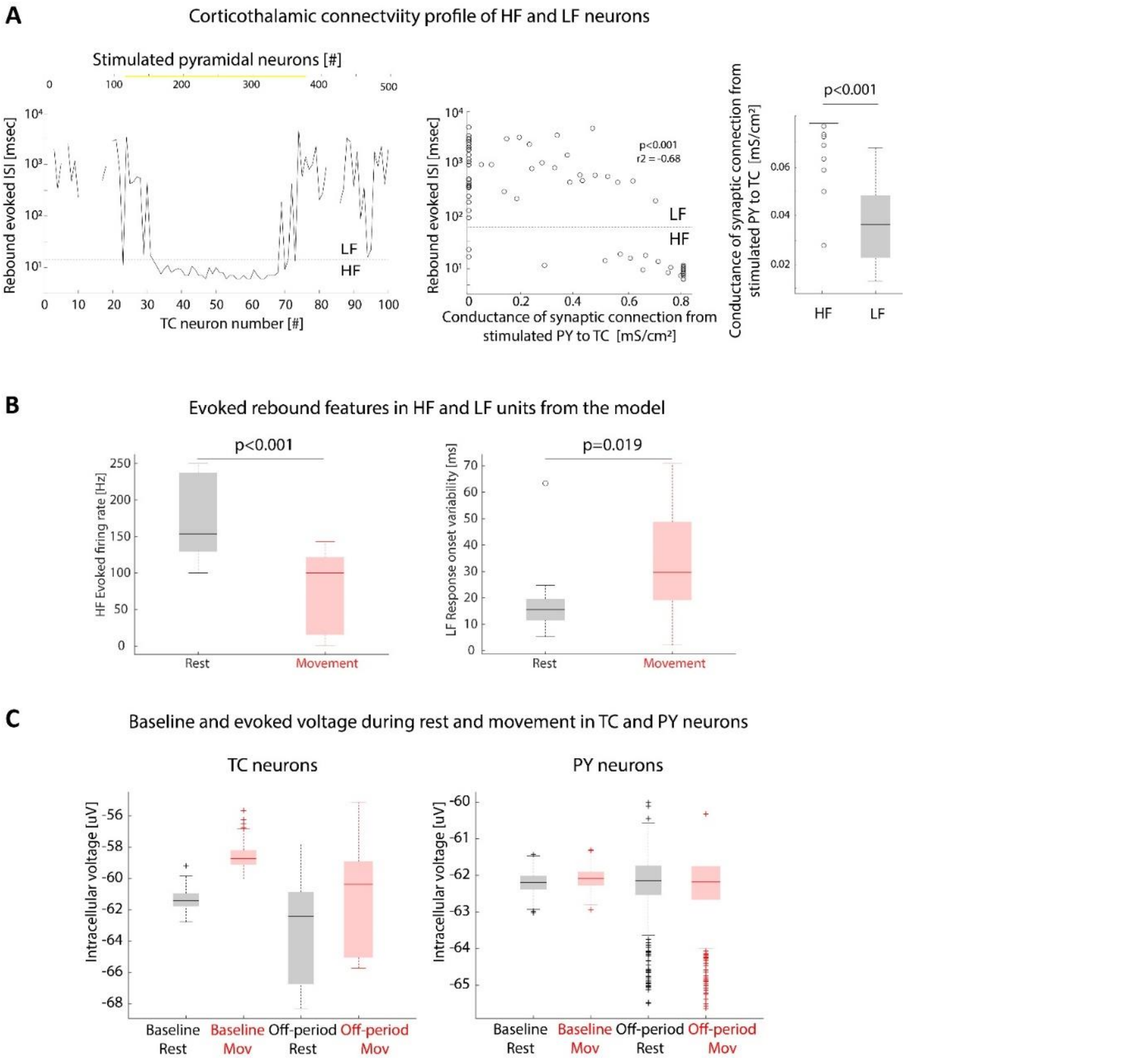


**Figure S14. Characterization of the cortical and thalamic responses to cortical stimulation in the simulated data.**

A) Characterization of the responses of TC neurons as a function of the conductance of the synaptic connections received by the stimulated PY neurons. Left: representation of the stimulated PY neurons (yellow). Middle: evoked inter spike interval (ISI) in the rebound response (75-250 ms) for each TC neuron. The dashed black line shows the threshold used to separate HF and LF neurons for the in-vivo results (17.88 ms, same as Figure 3C). Middle: evoked rebound ISI (75-250 ms) for each TC neuron as a function of synaptic input from the stimulated PY neurons (Pearson's correlation; $p<0.001$; $r=-0.68$). Right: synaptic inputs comparison between HF and LF neurons from the stimulated PY neurons. This quantification shows that HF receives significantly larger synaptic inputs from the stimulated cortical neurons compared to LF neurons (HF: 0.77±0. 098 uS; LF: 0.37±0. 16 uS; Wilcoxon rank sum test, $p<0.001$).

B) Difference between the thalamic response features evoked during rest and movement in the simulated data. Left: evoked firing rate for HF units (Wilcoxon Rank sum test; $p<0.001$). Right: response onset variability for LF units (Wilcoxon Rank sum test; $p=0.019$).

C) Median intracellular voltage during baseline and during 25-100 ms (off-period) response window for TC neurons (left) and PY neurons (right) during rest and movement.

| Sex | Handedness | Age |
|---|---|---|
| M | Left | 27 |
| F | Right | 61 |
| M | Right | 26 |
| M | Right | 38 |
| M | Right | 28 |
| M | Right | 25 |
| F | Right | 27 |
| M | Right | 48 |
| F | Right | 25 |
| F | Right | 25 |
| F | Right | 31 |
| M | Left | 31 |

**Supplementary Table 1. Demographics of the healthy subjects undergoing the TMS-EEG protocol.**

| Sex | Handedness | Age | Epileptic zone | iEEG hemisphere | Etiology | MRI | Drugs |
|---|---|---|---|---|---|---|---|
| M | Right | 29 | Left operculum and insula | Left | Gliosis | Negative | Oxcarbamazepine 1800 mg/die; Lacosamide 400 mg/die; Topiramate 300 mg/die; Clobazam 10 mg/die |
| M | Right | 44 | Left superior parietal lobule | Left | Focal cortical dysplasia IIa | Negative | Lacosamide 400 mg/die; Topiramate 400 mg/die; Brivaracetam 200 mg/die |
| F | Right | 27 | Right dorsal medial frontal lobe | Bilateral | Gliosis | Negative | Felbamate 1800 mg/die; Lamotrigine 400 mg/die; Carbamazepine 600 mg/die; Fenobarbital 100 mg/die |
| F | Right | 27 | Right insula | Right | Gliosis | Right fronto-central-opercular malformative lesion | Levetiracetam 2000 mg/die; Clobazam 10 mg/die; |
| M | Left | 23 | Frontal medial and frontal premotor left | Left | Unknown | Softening focus in the inferior central gyrus and hemispheric asymmetry | Carbamazepine 800 mg/die; Perampanel 6 mg/die; |

**Supplementary Table 2. Demographics and clinical information of the patients with epilepsy who underwent the ES-iEEG protocol.**
